# Who is the best surrogate for germ stem cell transplantation in fish?

**DOI:** 10.1101/2021.08.09.455047

**Authors:** Roman Franěk, Yu Cheng, Michaela Fučíková, Vojtěch Kašpar, Xuan Xie, Mujahid Ali Shah, Otomar Linhart, Ivo Šauman, Martin Pšenička

## Abstract

Surrogate reproduction technology in fish has potential for aquaculture as well as endangered species preservation and propagation. Species with some unfavourable biological characteristics for culturing such as a late maturation or a large body size are ideal candidates for surrogate reproduction using smaller and faster-maturing host. One of the general prerequisites for the successful surrogacy and the pure donor-derived gamete production is the sterility of the host. Various sterilization methods have been developed and used in fish surrogacy; however, a direct comparison of available methods is missing. Such a knowledge gap hinders choice for the surrogate in various fish species, including those in high commercial demand such as tuna or sturgeons, where is a particular limitation from the point of the live material availability and difficulty to perform a high throughput assessment of different surrogates. Yet, large sturgeons or tuna species are one of the most prominent candidates for surrogacy. Zebrafish was utilized in this study as a model species to answer whether and to which extent different sterilization strategies can affect the surrogacy. Germ cell-depleted recipients (produced using knockdown of *dead end* gene), triploid recipients, and zebrafish x pearl danio hybrid recipients were tested as they represent the most frequently used types of surrogates. Spermatogonia isolated from vas::EGFP transgenic strain were intraperitoneally transplanted into swim-up 5-day old zebrafish. Transplantation success, survival, gonadal development, and reproductive output of the fish was analyzed. Germ cell-depleted recipients with empty gonads were identified as the most convenient among tested sterilization methods considering surrogacy induction success and reproductive output. The present study stands as significant aid for selecting suitable surrogates in various fish species.

## 1 INTRODUCTION

Germ stem cell (GSC) manipulation in fish is still a relatively novel reproductive biotechnology. The stem potential of GSCs in gonads is used for surrogate production of donor-derived gametes. Isolated GSCs from an individual are transplanted into another individual, even from a relatively distinct species. Transplanted GSCs are capable of migration and genital ridge colonization. Afterwards, GSCs can undergo trans-differentiation when spermatogonia in a female body environment switch to an oogonial fate and vice versa. Transplanted GSCs can proceed with gametogenesis and give rise to donor-derived gametes (Goto and Saito, 2019). Surrogacy can be accompanied by the inclusion of cryopreservation procedures when both male (Franěk et al., 2019a) and female (Franěk et al., 2019b) GSCs can be cryopreserved efficiently and then recovered by transplantation into surrogate hosts (Lee et al., 2013; Yoshizaki and Lee, 2018). Moreover, GSCs manipulation technology in fish is recently being applied to produce genetically edited donor gametes while avoiding eventual inviability of adult individuals because of induced mutation (Zhang et al., 2021, 2020).

It is reasonable to presume that surrogate reproduction will be applied for species preservation or aquaculture since the number of reports on GSCs manipulation in various species increases rapidly, including aquaculture relevant species (Goto and Saito, 2019). Gametogenesis of large and late-maturing species might be accelerated by GSCs transplantation into smaller and faster-maturing recipients (Linhartová et al., 2015; Hamasaki et al., 2017; Baloch et al., 2019b). Reversely, surrogacy can be utilized to increase the gamete production by transplantation of GSCs from smaller to larger and potentially more fecund species, e.g. from goldfish (*Carassius auratus*) to common carp (*Cyprinus carpio*) or from sterlet sturgeon (*Acipenser ruthenus*) to beluga (*Huso huso*). Surrogacy also has potential to ameliorate breeding schemes via the distribution of the germplasm from superior individuals via surrogates (Jin et al., 2021; Yang et al., 2021; Yoshizaki and Yazawa, 2019).

Since surrogate reproduction in fish is a long term and laborious effort, whole technology needs to be optimized to maximize its success. Optimal conditions for GSCs isolation (Shikina et al., 2013), purification (Ryu and Gong, 2020), and *in vitro* expansion (Iwasaki-Takahashi et al., 2020; Xie et al., 2019) were identified. Several studies already paid attention to the investigation of variables related to used recipients. Optimal developmental stages for donors and recipients were investigated in salmonids by primordial germ cells (PGCs) transplantation (Takeuchi et al., 2003) and in zebrafish (*Danio rerio*) single PGCs transplantation into the blastula stage host. PGCs were shown to lose their migratory potential progressively in zebrafish (Kawakami et al., 2010; Saito et al., 2010). Similarly, differences were demonstrated on medaka (*Oryzias latipes*) in the age of recipients for spermatogonia transplantation (decreasing transplantation success with increasing age) and in the positive effect of higher number of transplanted spermatogonia on the colonization rate (Seki et al., 2017). Also, a short time window for transplanted GSCs to incorporate into the host’s genital ridge has been identified in rosy bitterling (*Rhodeus ocellatus*), suggesting certain biological limitations of the transplantation procedure (Octavera and Yoshizaki, 2018).

However, many other factors affecting surrogacy success have been described initially, such as the potential influence of the genetic relativeness between donor and recipient on the transplantation success (Takeuchi et al., 2003). Or a comparison of the sterile and unsterile recipient (Marinović et al., 2019) or a consequence of different sterilization methods on surrogacy success (Octavera and Yoshizaki, 2018). Sterilization is crucial for successful surrogacy since introduced GSCs do not have to compete for gonadal niche, and adult surrogates can produce only donor-derived gametes. However, unlike in mammals, fish can be sterilized by diverse ways and mechanisms, resulting in various levels of sterility ranging from germ cell-free gonads to decently developed gonads incapable of producing motile spermatozoa.

Available methods for surrogate larvae sterilization are based on complete PGCs depletion via targeting *dead end* (*dnd*) gene necessary for PGCs migration and maintenance (Baloch et al., 2019a). PGCs depletion can be achieved by a temporal inhibition of translation using antisense morpholino oligonucleotide proven to be effective in several fish species – sterlet sturgeon (*Acipenser ruthenus*) (Linhartová et al., 2015), loach (*Misgurnus anguillicaudatus*) (Fujimoto et al., 2010), goldfish (*Carassius auratus*) (Goto et al., 2012), cod (*Gadus morhua*) (Škugor et al., 2014), zebrafish (Slanchev et al., 2005), rainbow trout (*Oncorhynchus mykiss*) (Yoshizaki et al., 2016). More recently, gene editing methods such as CRISPR/Cas9 or zinc finger nucleases have been employed to target *dnd* gene in sterlet sturgeon (Baloch et al., 2019b), Atlantic salmon (*Salmo salar*) (Wargelius et al., 2016) and zebrafish (Li et al., 2017). Result of this sterilization method are gonads utterly free of the GCs; however, the nature of this sterilization method can be considered challenging and laborious since it requires individual embryo injection and known *dnd* sequence. It is also necessary to be aware that sterilization by gene editing is considered as genetic modification which might result in more strict regulations on maintenance and use of genetically modified fish.

Triploidization is another method of choice for sterility induction. Triploids are produced by chromosome manipulation via the second polar body retention by a shock briefly after the fertilization. Artificially induced triploids usually have impaired gametogenesis as a consequence of odd chromosome number resulting in synapsis defects during meiosis (Piferrer et al., 2009). Protocols for triploidy induction have been developed in many species (Piferrer et al., 2009), some of them were also used as surrogates and successful donor-derived gametes were finally produced from triploid rainbow trout (Lee et al., 2013), Atlantic salmon (Hattori et al., 2019), medaka (Seki et al., 2017) and zebrafish (Franěk et al., 2019c) recipients. Triploidy is suitable for large scale production of recipients. However, there are also species for which triploidy does not guarantee complete sterility (Murray et al., 2018), thus triploid recipients need to be used with cautions.

Last practically feasible method for sterilization for before intraperitoneal transplantation is an interspecific hybridization. Fishes are, in most cases, external fertilizers which enable their simple production, including hybridization. Hybridization has been attempted in many species, often resulting in impaired reproductive performance. Reasons for altered gonadal development are conditioned by genetic (in)compatibilities of two parental species (Fujimoto et al., 2008; Tichopád et al., 2020), such as altered epistasis (Orr and Irving, 2001). Gonad in zebrafish (ZF) x pearl danio (PD) (*Danio albolineatus*) hybrids can develop into male or female-like structures (Wong et al., 2011). The presence of both sexes was also observed in the hybrid of two marine species, blue drum (*Nibea mitsukurii*) and white croaker (*Pennahia argentata*), reporting arrested PGCs not proliferating further (Yoshikawa et al., 2018). Both studies also tested infertile hybrids as surrogates. They confirmed successful donor-derived gamete production showing that hybrid sterility is GSC autonomous when supportive gonadal somatic cells are likely to remain unimpaired by hybridization and can nurse transplanted GSCs. Hybridization among different tetra species resulted in various patterns of gonadal development. Usually, diploid hybrids possessed gonads with distinguishable male or female phenotypes with few germ cells; however, even advanced stages of gonadal development might occur (Piva et al., 2018). Thus, hybridization is not a universal approach for sterile surrogate production, and careful evaluation must be done in advance. On the other hand, hybridization is suitable to facilitate large scale production of recipients.

There are several other methods for sterilization before germ cell transplantation, such as a combination of thermal and cytostatic treatment. These methods are rather suited for intrapapillary GSCs transplantations conducted in adult or juvenile fish (Lacerda et al., 2010; Nóbrega et al., 2010). Similarly, sterilization is possible using specific transgenic lines by interfering PGCs migration via ubiquitous expression of SDF1 (essential for PGCs migration) triggered by thermal treatment (Wong and Collodi, 2013) or with nitroreductase expression in PGCs exclusively by immersion into metronidazole enzyme resulting in its conversion into toxic metabolites targeting only PGCs in zebrafish causing their depletion (Zhou et al., 2018). Transgenic medaka strain with follicle-stimulating receptor mutation causing sterility in females was sex-reversed into phenotypic males with subsequent spermatogonia transplantation into sterile hybrids of *Oryzias latipes* and *O. curvinotus* to rescue egg production while maintaining the mutation transmission. Subsequently, sperm from sex-reversed females homozygous for follicle-stimulating receptor mutation was used, and a system for production of all-female sterile progeny was established (Nagasawa et al., 2019). All the above-mentioned transgenic strategies for sterilization are compelling and effective once given transgenic line is established. Unfortunately, not all species are convenient for transgenesis due to their long generation times and the lack of genomic data.

We presume that *dnd* knockdown, triploidization and hybridization are the most practical and universal methods to sterilize recipients before the intraperitoneal germ cell transplantation. Theoretically, completely germ cell less gonads might represent the best environment for transplanted GSCs, as they do not need to compete for space with endogenous germ cells during colonization. However, exhaustive comparison of different sterilization methods has not been performed yet. The presented study aimed to provide a comprehensive analysis of different sterilization treatment and their effect on surrogacy success in fish.

## 2 Material and methods

The study was conducted at the Faculty of Fisheries and Protection of Waters (FFPW), University of South Bohemia in České Budějovice, Vodňany, Czech Republic. The facility has the competence to perform experiments on animals (Act no. 246/1992 Coll., ref. number 16OZ19179/2016–17214). The expert committee approved the methodological protocol of the current study of the Institutional Animal Care and Use Committee of the FFPW according to the law on the protection of animals against cruelty (reference number: MSMT-6406/119/2). The study did not involve endangered or protected species. Authors of the study (RF, MF, VK, OL, MP) own the Certificate of professional competence for designing experiments and experimental projects under Section 15d (3) of Act no. 246/1992 Coll. on the Protection of Animals against Cruelty.

### 2.1 Fish and production of recipients

Zebrafish broodstock was maintained as described previously (Franěk et al., 2019c). Zebrafish AB line (descendants of fish purchased from European Zebrafish Resource Centre), vas::EGFP line (EGFP expression is under control of vasa promotor) (descendants of fish purchased from University of Liège, Belgium) and pearl danio (descendants of fish purchased from PetraAqua, Czech Republic). All experimental groups were produced by *in vitro* fertilization. For each transplantation trial, pooled eggs and pooled sperm was divided into four groups to establish recipient groups from the same parents. PGCs depleted fish by *dnd*-MO (MO group) were produced by injecting zebrafish embryos at 2-8 cell stage with 100 µM solution of antisense morpholino diluted in 0.2 KCl with 1.5% Rhodamine B isothiocyanate–Dextran (10,000 MW) to ease the confirmation of successful MO injection by fluorescence signal detection in the animal pole. To produce triploids (3n group), an optimized heat-shock protocol (Franěk et al., 2019c) using 41.4 °C, initiated 2 min post fertilization (mpf), lasting 2 min was performed. Hybrids between zebrafish females and pearl danio males (H group) were produced as described previously (Wong et al., 2011). Recipients were produced in three replicates at different timepoints (Fig. 1). Produced embryos were cultured at 28.5 °C, swim-up larvae were fed from 5^th^-day post fertilization (dpf) with paramecium *ad libitum*, from 10 dpf with *Artemia* sp. At the age of 4 weeks post-fertilization (wpf) fish were transferred into zebrafish housing system and fed with a combination of dry diet and *Artemia* sp. until the termination of the experiment.

**Figure 1.**
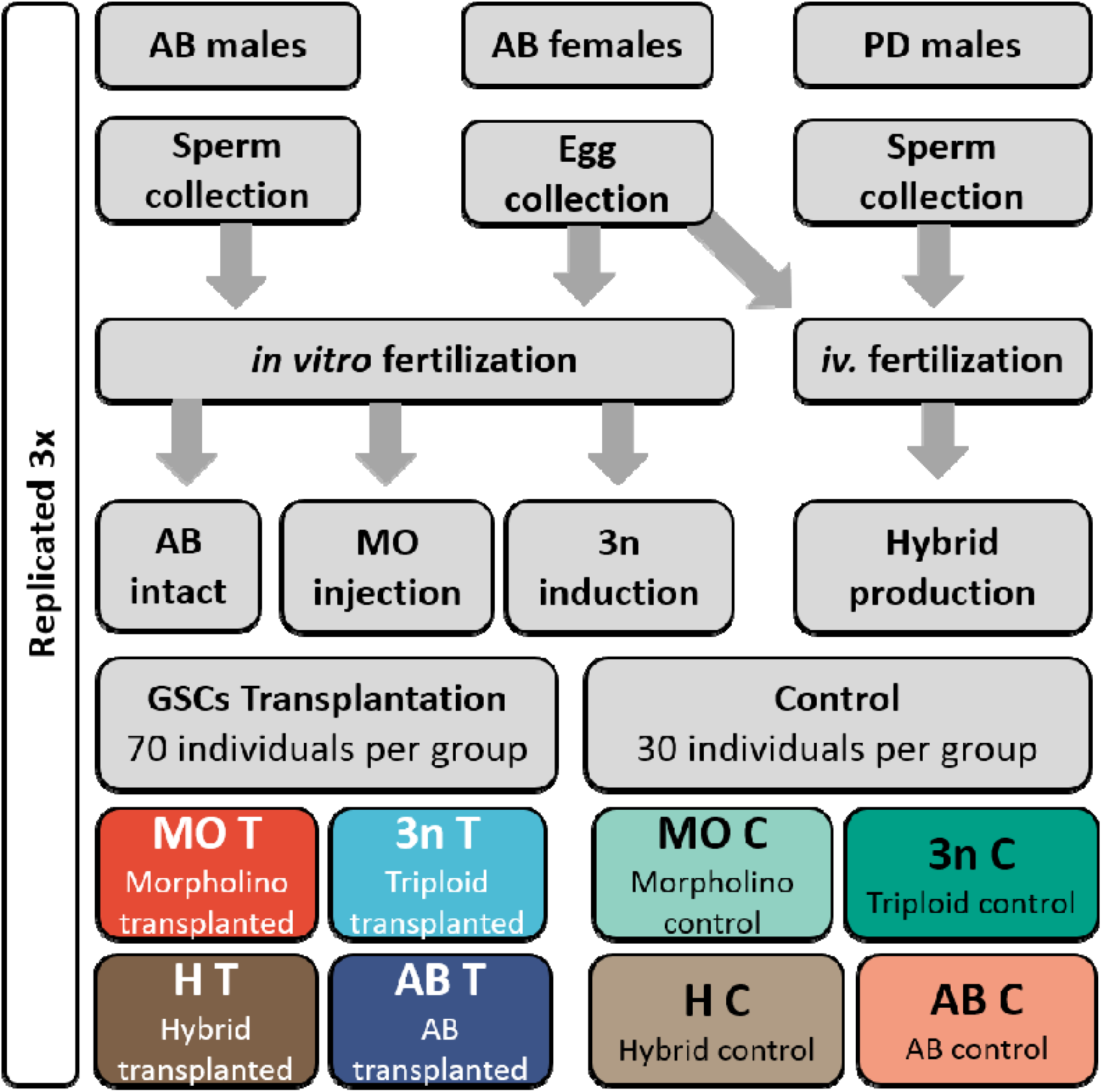
Production of experimental and control groups.

### 2.2 Germ cell donors and transplantation

Adult zebrafish donor males (6-8-month-old) from vas::EGFP transgenic line were over anaesthetized in MS222, body was washed with 70% ethanol and decapitated. Testes were removed carefully (8 males for one transplantation trial) and kept in phosphate-buffered saline (PBS). Testis were cut into small fragments in 2 ml tube with 0.1 ml of PBS and washed thoroughly by several changes of PBS to facilitate sperm leakage. Afterwards, finely cut testes were digested in 8 ml of dissociation media containing 0.1% trypsin in PBS on a laboratory shaker at room temperature for 60 min. During digestion, 0.5% DNase solution in distilled water was added when clumping or agglutination of testis fragments was observed (usually, 70-100 µl of DNase was used). Digestion was terminated by the addition of 7 ml L15 media with 20% fetal bovine serum (FBS). The suspension was filtrated through a sterile 30µm mesh filter, centrifuged at 0.4 g for 10 min. The supernatant was removed, and the pellet was resuspended in fresh L15 with 10% FBS and stored at 10 °C during transplantation.

Transplantation was performed at 5 dpf. Recipients were anaesthetized in 0.05% MS222 and placed on agar coated petri dish. Microcapillary was polished on a grinder, filled with cell suspension and mounted on micromanipulator with a pneumatic injector to keep equal injection pressure and duration during transplantation. Seventy fish from each recipients group were transplanted in three replicates. Always 10 recipients per group were transplanted, and then 10 recipients from another group were transplanted to minimize the chance that some groups would be transplanted with “aged” isolated testicular cells. Four transplanted groups were established – MO T, 3n T H T and AB T. Controls were established from non-transplanted fish, 30 specimens for each recipient group and replicate – MO C, 3n C, H C and AB C.

### 2.3. Identification of germline chimeras

Two weeks post-transplantation all surviving fish from transplanted groups were anaesthetized and screened under a fluorescent stereomicroscope (Leica M205 FA) and separated to fish with EGFP signal positive and negative. EGFP positive fish were evaluated for EGFP positive transplanted cells distribution. All surviving transplanted adult fish from EGFP positive groups were prepared for sperm collection as described previously (Franěk et al., 2019c). Sperm was collected individually into 20 μl of E400 media (Cheng et al., 2021) and observed under a fluorescent microscope (Olympus IX 83) to detect EGFP signal witnessing donor-derived origin. The remainder of collected sperm was used for genotyping. DNA was extracted using the Hot-Shot method and PCR with EGFP specific primers followed by gel electrophoresis as described previously (Franěk et al., 2019c). Afterwards, confirmed chimeric males were reared separately from males not producing donor-derived sperm.

### 2.4 Reproductive performance of germline chimeras

#### 2.4.1 Fertilization tests

Confirmed chimeric males were set up randomly into spawning aquaria (1l) with females from control AB strain (1:1) as described for semi-artificial spawning (Franěk et al., 2019c) and allowed to spawn for 4 hours. Eggs were collected, and their survival was monitored (fertilization rate, 24 hours post-fertilization (hpf), 48 hpf, hatching rate and swim-up rate). After hatching, 10 larvae from each cross were frozen fixed for genotyping with EGFP specific primers. For each recipient group, data from 15 successful spawnings were collected for assessment of spawning success, and number of oviposited eggs (note that individual males were not spawned repeatedly) were collected. For *in-vitro* fertilization, sperm from three males (note that males were not used repeatedly) from each experimental group, including controls (also vas::EGFP strain), was collected individually, then pooled and used to fertilize fraction from the mixture of stripped eggs from AB control females. In total, *in-vitro* fertilization was repeated three times (with different males).

Females from AB T were firstly anaesthetized and screened under a fluorescent steromicroscope to detect the presence of EGFP signal in the ovaries. EGFP positive AB T females were then attempted to be spawned semi-artificially, collected eggs were separated according to EGFP signal, and their fertilization and hatching rates were monitored.

#### 2.4.2 Spermatozoon motility, velocity and sperm concentration assessment

Sperm samples collected from anaesthetized fish were stored immediately into 20 μl E400. Distilled water containing 0.25% Pluronic F-127 (to prevent spermatozoon from adhering to microscope slides) was used as the activation medium. Activation medium and sperm samples were stored on ice before motility activation. Sperm was activated at room temperature (21 °C) by mixing the immobilized sperm sample into 20 µl of the activation medium on a glass slide within 1 hour post collection. The activated spermatozoa were directly recorded microscopically (UB 200i, PROISER, Spain) at 10× using a negative phase-contrast condenser with an ISAS digital camera (PROISER, Spain) setting at 25 frames/s. The Integrated System performed analyses of the sperm recordings for Sperm Analysis software (PROISER, Spain) at 15 s post sperm activation. Computer-assisted sperm analysis included the percentage of motile sperm (%), curvilinear velocity (VCL, μm/s), straight-line velocity (VSL, µm/s), and spermatozoa rate with rapid motility (> 100 μm/s), medium motility (46 to 100 μm/s), slow motility (10 to 45 μm/s), and static spermatozoa (< 10 μm/s). Analyses of all samples (9 males per recipient group) were carried out in triplicate (each male recorded three times).

Sperm concentration and the total number of sperm per male were evaluated for individual males in E400 extender solutions. The sperm in E400 was diluted again 10-140 times according to the density of sperm. The sperm concentration (expressed as 10^6^/µl) was determined by a Bürker cell haemocytometer (Marienfeld, Germany, 12 squares counted for each male) using an optical phase-contrast condenser and an ISAS digital camera (PROISER, Spain) under an Olympus microscope BX 41 (4009). All measurements were repeated 3 times.

### 2.5 Histology

Sacrificed fish were firstly degutted, decapitated and photographed under a fluorescent microscope. Trimmed torsos with gonads inside were fixed in Bouin’s fixative overnight, washed in 70% ethanol, and processed by resin sectioning and haematoxylin-eosin staining (Sullivan-Brown et al., 2011). At least three specimens from transplanted and non-transplanted groups, including controls, were processed.

### 2.6 Electron microscopy

Sperm collected in E400 media (from AB C and H C group) was fixed in 2.5% glutaraldehyde in PBS. Samples were prepared for electron scanning microscopy as described previously (Franěk et al., 2021) and observed on JEOL JSM-7401F scanning electron microscope.

### 2.7 Confocal microscopy examination

Dissected gonads were fixed in 4% paraformaldehyde (PFA) in PBS for 2h, washed 3 times in PBS and immersed for 3h in 25% sucrose solution (in PBS). Specimens were incubated overnight in Cryomount media (HistoLab), then placed and oriented in fresh Cryomount media into plastic moulds and frozen on floating Styrofoam (1 cm height) in liquid nitrogen vapours and stored at - 80 °C until use. Frozen tissue blocks were equilibrated in the cryostat chamber at -25 °C for 30 min before cutting and attached to metal chucks. Sections of 15 µm thickness were cut on cryostat, attached on superfrost slides, allowed do dry at RT for 5 min and mounted in Fluoroshield™ with DAPI histology mounting medium, sealed with coverslip and imaged with laser scanning confocal microscope (Olympus FV 3000).

### 2.8 Data evaluation and statistical analysis

The data homogeneity of dispersion was evaluated using Levene’s test. The difference in sperm motility parameters, survival, 24 and 48 hours post-fertilization (hpf), hatching, swim-up among groups was analyzed using a one-way ANOVA. LSD test determined all the differences among means. Bar charts of survival rates in fertilization, 24 and 48 hpf, hatching and swim-up, survival percentage in transplanted, 1, 7 and 14 days post-transplantation (dpt), 1, 3 and 6 months post-transplantation (mpt), EGFP positive cells location (%) were drawn with mean ± standard deviation of the mean (S.D.). All analyses were performed at a significant level of 0.05 by using R (R Core Team, 2018).

## 3 RESULTS

### 3.1 Survival, transplantation success and colonization patterns

The lowest survival during recipient production was observed in MO and 3n group which is attributed to injection into embryos and heat shock treatment respectively. In hybrid group we observed increased mortality prior to hatching (Fig. 2A). However, given the nature and easiness of zebrafish breeding, we do not consider lower survival due to sterilization treatment as limiting. Notably, post-transplantation survival in MO T group was comparable to the controls, while survival performance of 3n T, H T and control groups was slightly lower. Altogether, overall survival from transplantation to 6 months of age was in all groups (included transplanted groups) from 65 to 85 % (Fig. 2B).

**Figure 2.**
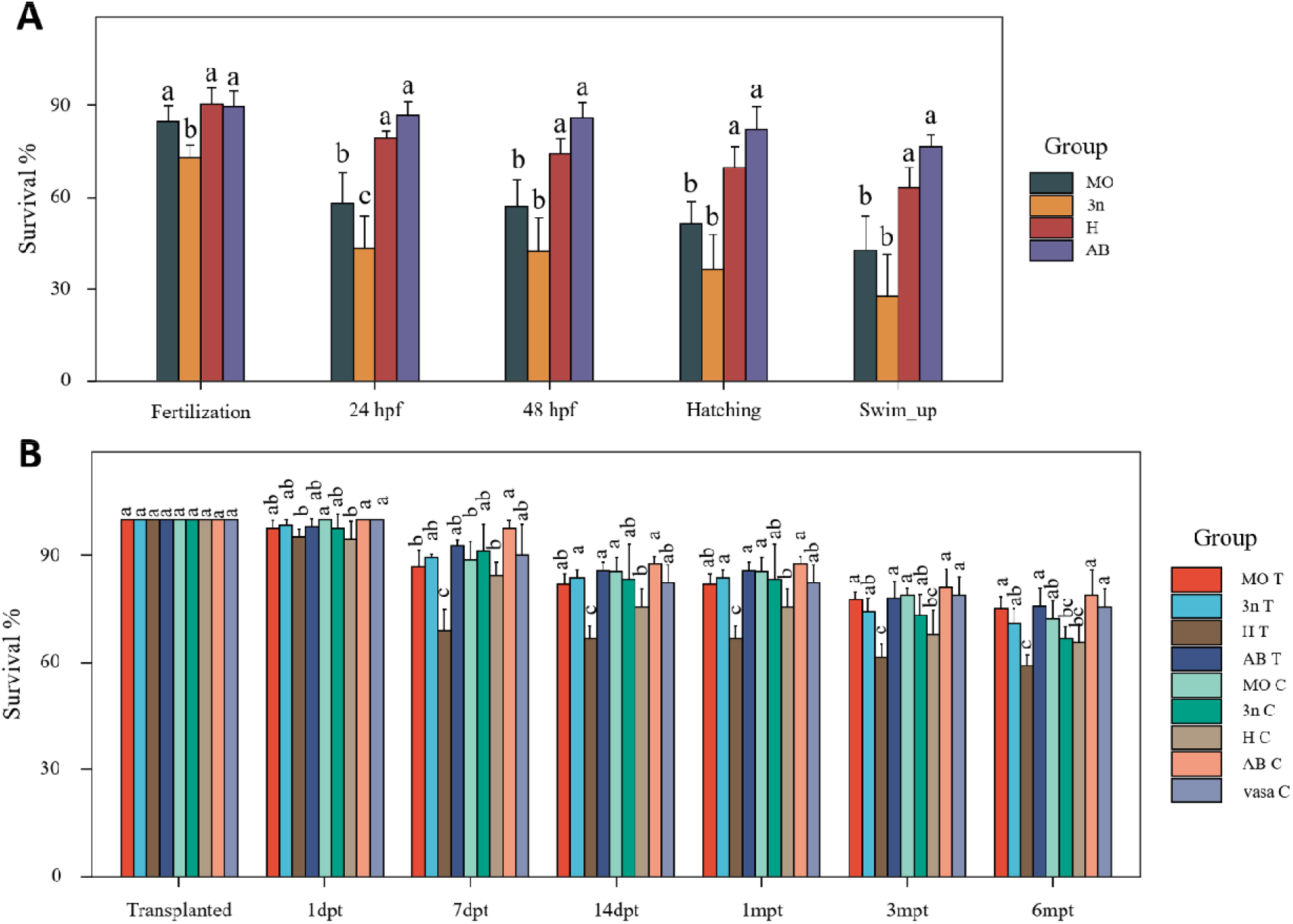
Overall survival after different sterilization treatments. Survival from fertilization to swim-up stage (A) and survival from transplantation to 6 months of age (B). (mean ± S.D.). Different letter denotes statistically significant difference between groups at each developmental stage (P < 0.05, one-way analysis of variance (ANOVA) followed by an LSD test for post hoc multiple comparisons).

The transplantation success evaluated two-week post-transplantation showed consistent results across different sterilization methods of the EGFP positive cells in recipients (Fig. 3A). Most of the EGFP positive cells were located in the posterior or medial part of the body cavity (Fig. 3B1-2). The anterior part was occupied by the EGFP positive cells rarely. This trend was prominent primarily in MO and H recipients, while 3n and AB recipients showed more similar colonization patterns between the posterior and medial part of the body cavity.

**Figure 3.**
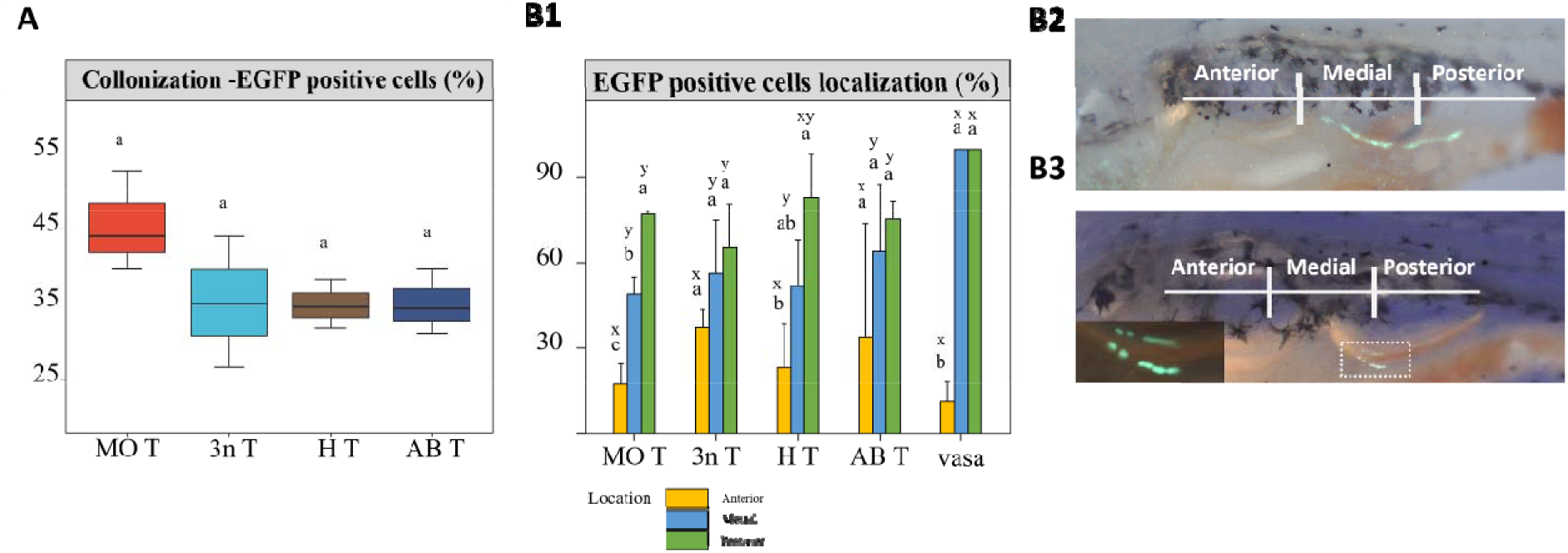
Colonization and cell localization after vas::EGFP GSCs transplantation. A) Comparison of transplantation success evaluated in all recipient groups at 2 wpt. No significant differences were detected among groups. B1) Localization patterns of EGFP positive cells in the body cavity of the recipients. Different superscripts, i.e. a, b, and c, indicate statistical differences between location for the same group whereas different superscripts, i.e. x and y, indicate statistical differences between the group at the same location (P < 0.05, one-way analysis of variance (ANOVA) followed by an LSD test for post hoc multiple comparisons). B2, B3) Evaluation of EGFP positive cells localization in the body cavity. B2) vas:EGFP control individual. B3) MO T individual. Cells in the white dashed rectangle are example of medial-posterior colonization.

### 3.2 Gonadal development in sterilized controls and surrogates

Sterilization treatments resulted in distinct patterns of gonadal development of adults. Testes of MO C treated fish were small and free of germ cells. Stromal somatic cells formed cavities divided by connective tissue into smaller compartments (Fig. 4A). In MO C group all (n=10) assessed controls showed germ cell-free testis. Control triploids developed gonads with all spermatogenic stages but with apparent defects in meiosis, resulting in aberrant spermatogenesis and the presence of only few spermatozoa in testicular lumens (Fig. 4B). Gross appearance of 3n testis was smaller than normal testis of diploid, while being more transparent due to lack of the high number of spermatozoa inside (Fig. 4B2). Sections from triploids showed consistent testicular development in all assessed specimens (n=10). More erratic development was observed in H C group. Gonads of hybrid males exhibited three phenotypes. H C had well-developed testis with few abnormally sized spermatozoa (H I.) (Fig. 4C), nearly one third of hybrids had undeveloped testis lacking germ cells (H II.) (Fig. 4D) and few individuals showed combination of one developed and one undeveloped testis (H III.) (Fig. 4E). Spermatozoa in developed hybrid testis were apparently large, which was later observed with light microscopy. Gonadal phenotype in hybrids was presented mainly by H type I. (well-developed gonads), H type II. males were less abundant, and only few individual fish were identified as hybrid females (Fig. 5B5). Hybrid females developed gonads in decent size and were determined by inner lamella-like structure forming empty cavities. Details on hybrid spermatozoa, histology and incidence of gonadal phenotypes are given in figure 5. Hybrid ovaries were mostly composed of homogenous mass of the larger cells suggesting oogonia or early-stage meiotic oocytes (Fig. 4F). The incidence of hybrid females was rare and only 6 females were detected from all adult surviving hybrids controls (N=59). Detailed description of gonadal phenotypes from histological sections is given in figure 4.

**Figure 4.**
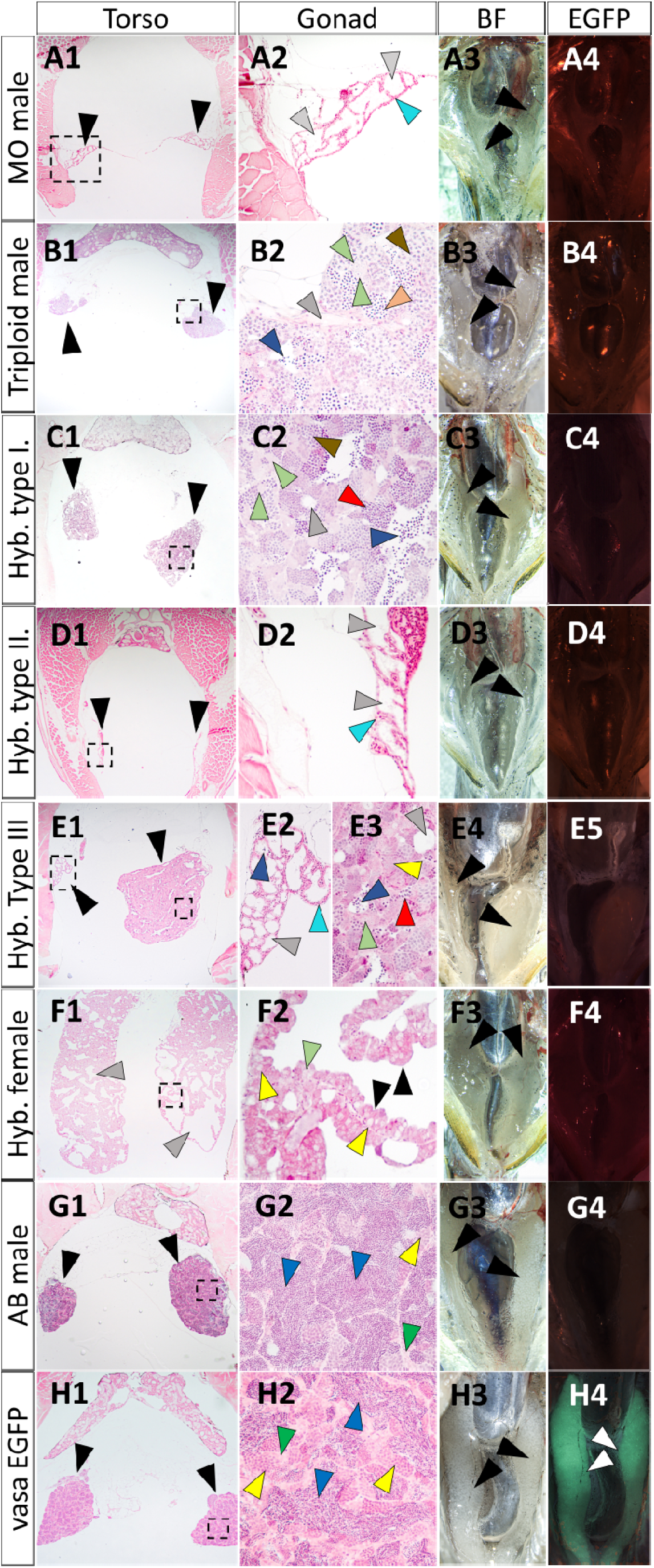
Different sterilization treatments affect gonadal development. **A) Morpholino treated male.** Both gonads are developed as empty testes (A1 – black arrowheads) lacking germ cells only from stromal cells (A2 – turquoise arrowhead) forming empty lumen-like structure (A2 – grey arrowheads). Gonads are macroscopically thin (A3 – black arrowheads) and without EGFP expression (A4). **B) Triploid male** with developed testis (B1 – black arrowheads). Only few individual spermatozoa are present in lumens (B2 – blue arrowheads), most of the lumens are sperm free and undeveloped (B2 – grey arrowhead). Meiotic germ cell arrested pachytene (B2 – brown arrowhead) and spermatids (B2 – orange arrowhead) are frequently observed as well as early-stage germ cells (B2 – green arrowheads). Gonads are macroscopically well developed, but without typical white colour (B3), lacking EGFP expression (B4). **C) Hybrid male with type I. gonads**. Testes are well developed (C1 - black arrowheads). Lumens are large with several dozens of spermatozoa with various head size (C2 – blue arrowhead), several empty and small lumens are present as well (C2 – grey arrowheads). Spermatocysts have clear structure and are mostly filled with meiotic germ cells with regular morphology (C2 – brown arrowhead) and with meiotic germ cells showing aberrant nuclei morphology (C2 – red arrowhead). Early-stage germ cells are frequently observed (C2 – green arrowheads). Gonads are macroscopically well developed (C3 – black arrowheads) and lacking EGFP signal (C4). **D) Hybrid male with type II. gonads – fully sterile**. Testes are undeveloped (D1 – black arrowheads) with similar morphology to MO treated male. Only stromal cells are present (D2 – turquoise arrowhead) forming empty compartments lacking spermatozoa (D2 – grey arrowheads). Gonads are macroscopically undeveloped and thin (D3 – black arrowheads) and lacking EGFP signal (D4). **E) Hybrid male type III. gonads**. One testis is poorly developed with similar structure to type II gonads, but few spermatozoa are present (E2 – blue arrowhead). Second testis is over developed with same structure and macroscopical appearance as in hybrid gonad type I (see C2 and C3) EGFP signal is not detected (D4). **F) Hybrid female**. Both ovaries are large with several empty inner compartments (F1 – grey arrowheads). Ovarian cells are forming typical lamella-like structure (F2 – black arrowheads), only early meiotic oocytes (F2 – yellow arrowheads) and early-stage germ cells are present (F2 – green arrowhead). Macroscopic structure is similar to hybrid male type I. gonads (F3 – black arrowheads), no EGFP signal is detected (F4). **Control recipient AB male (G) and donor vas::EGFP male (H)**. Testes are well developed, lumens are densely filled with spermatozoa (G2, H2 – blue arrowheads), several spermatocysts with meiotic spermatocytes are detected on the sections (G2, H2 – yellow arrowheads) as well as early-stage germ cells (G2, H2 – green arrowheads). Gonads are macroscopically well developed with typical white compartments indicating presence of sperm in large amount (H3, G3 – black arrowheads). Strong EGFP signal is detected in vas::EGFP testis (H4 – white arrowheads).

**Figure 5.**
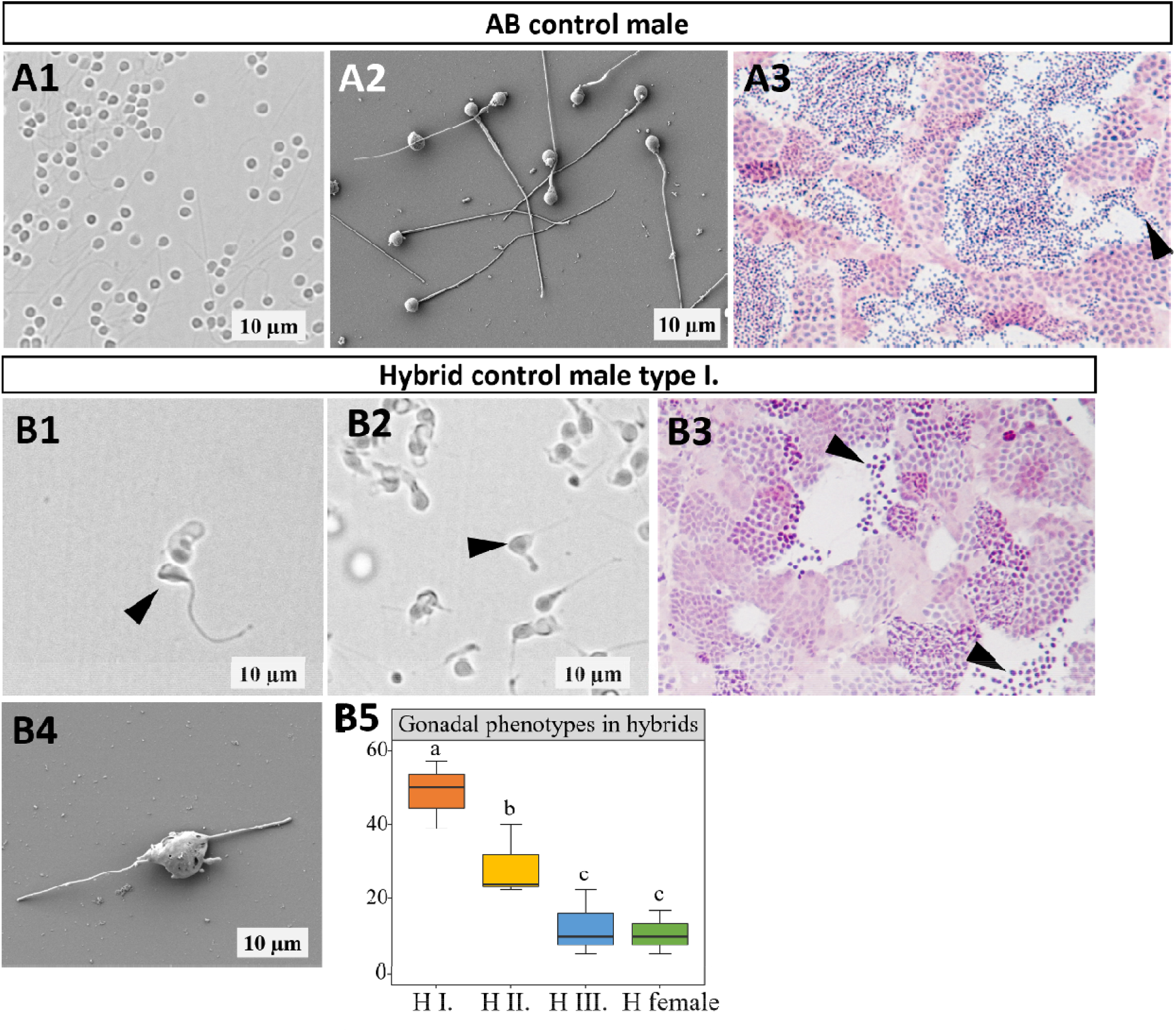
Abnormal spermatozoa morphology in ZF x PD hybrids. A) Spermatozoa appearance in AB control. A1) Light microscopy caption. A2) SEM caption showing normal spermatozoa morphology. A3) Traversal testicular section. B) Spermatozoa appearance in hybrid with gonadal type I. B1, B2) Light microscopy showing abnormally large and shaped heads of the spermatozoa, both flagellums are very short. B3) Transversal testicular section. Note the size of the cells in the lumen (black arrowhead) compared to size of cells in A3. B4) SEM caption of abnormal spermatozoa, with double head and two flagellums. B5) Percentage occurrence of different gonadal phenotypes (described in Fig. 4) in control hybrid group. Different letter denotes statistically significant difference (P < 0.05, one-way analysis of variance (ANOVA) followed by an LSD test for post hoc multiple comparisons).

### 3.3 Success in germline chimera male induction

The highest incidence of adult germline chimeras was observed in the MO T group, followed by 3n T group. Interestingly, % of adult germline chimeras in H T and AB T group were almost equal (Fig. 6A). Intraperitoneally transplanted GSCs were capable of establishing donor-derived spermatogenesis in all tested sterilization treatments as well in non-sterilized AB recipients. Dissection of adult germline chimeras showed influence of sterilization treatments on the extent of testicular development. Various patterns of gonadal development were observed across tested sterilization treatments and were reflected in GSI when compared to their respective controls (Fig. 6B). Observed gross gonadal development was prominent MO T group which gained the largest increment in gonadal development (comparing transplanted group with their respective sterilization control). In MO T group, transplanted GSCs were able to frequently reconstitute spermatogenesis unilaterally (Fig. 6 D3, 6 D4) or even bilaterally into fully developed testes in term of length and width (Fig. 6 D9). Similar capacity was also observed in 3n T and H T group, but with less incidence of fully developed gonad and higher incidence of only spatially localized EGFP positive spermatogenesis not reaching full length and width of the testes (Fig. 6D). Developed recipient-derived spermatogenesis in AB T group largely limited EGFP positive spermatogenesis when only spatially restricted EGFP positive areas of testicular tissue were observed macroscopically (Fig. 6 D5-6).

**Figure 6.**
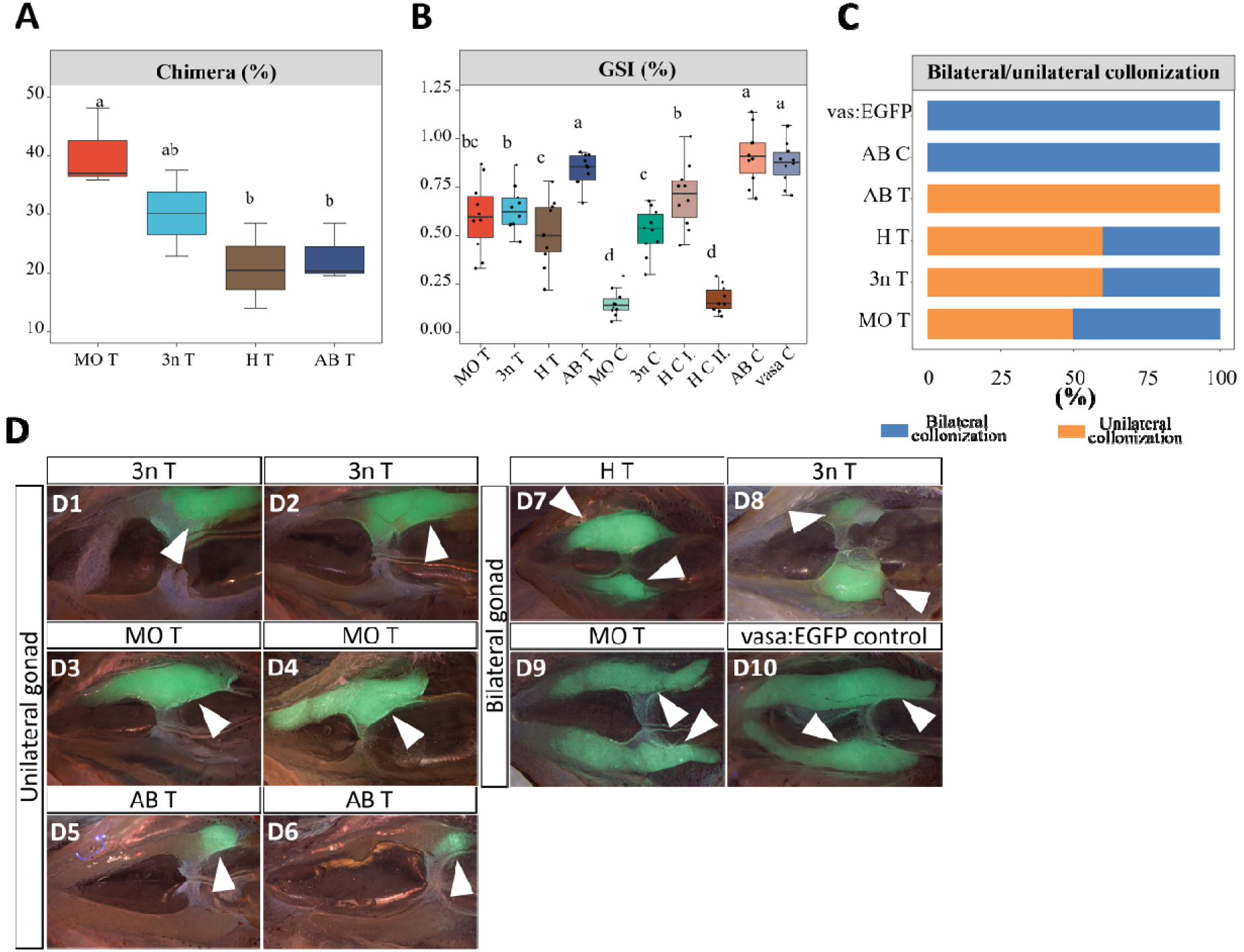
Adult germline chimeras. A) Overall % proportion of confirmed germline chimeras (in case of AB T also chimeric females were detected) (producing EGFP positive gametes) from surviving adults. B) GSI in experimental males and their respective control groups. C) Lateral patterns of testicular development in adult male chimeras. D) Ventral view on dissected germline chimeric male and donor control showing lateral and spatial patterns of testicular development. Values with different letters are significantly different (P < 0.05, one-way analysis of variance (ANOVA) followed by an LSD test for post hoc multiple comparisons).

Sterilization treatments promoted donor’s germ cells development. Introduced GSCs could expand and occupy whole testis cross-section, which was reflected by EGFP signal detection on cryosections (Fig. 7). Interestingly, triploid and non-sterilized recipients showed that introduced GSCs must compete for the testicular niches with endogenous germ cells since cryosections in germline chimeras showed presence of spermatocysts lacking EGFP signal (Fig. 7B). Germline chimeras from AB T group exhibited more erratic distribution of exogenous germ cells which was limited spatially and EGFP positive cells were not able to occupy the recipient testes completely (Fig. 7D-E). Histological analysis of identified germline chimeras showed testis with similar morphology to donor and recipient controls (Fig. 8). However, in some individuals from 3n T and H T group we have identified signs about partial spermatogenesis of recipient’s germ cells e.g. spermatocysts with meiotic germ cells arrested in pachytene (3n T) as well as individual spermatozoa with abnormally large heads and empty testicular lumens in H T group (Fig. 8 C4).

**Figure 7.**
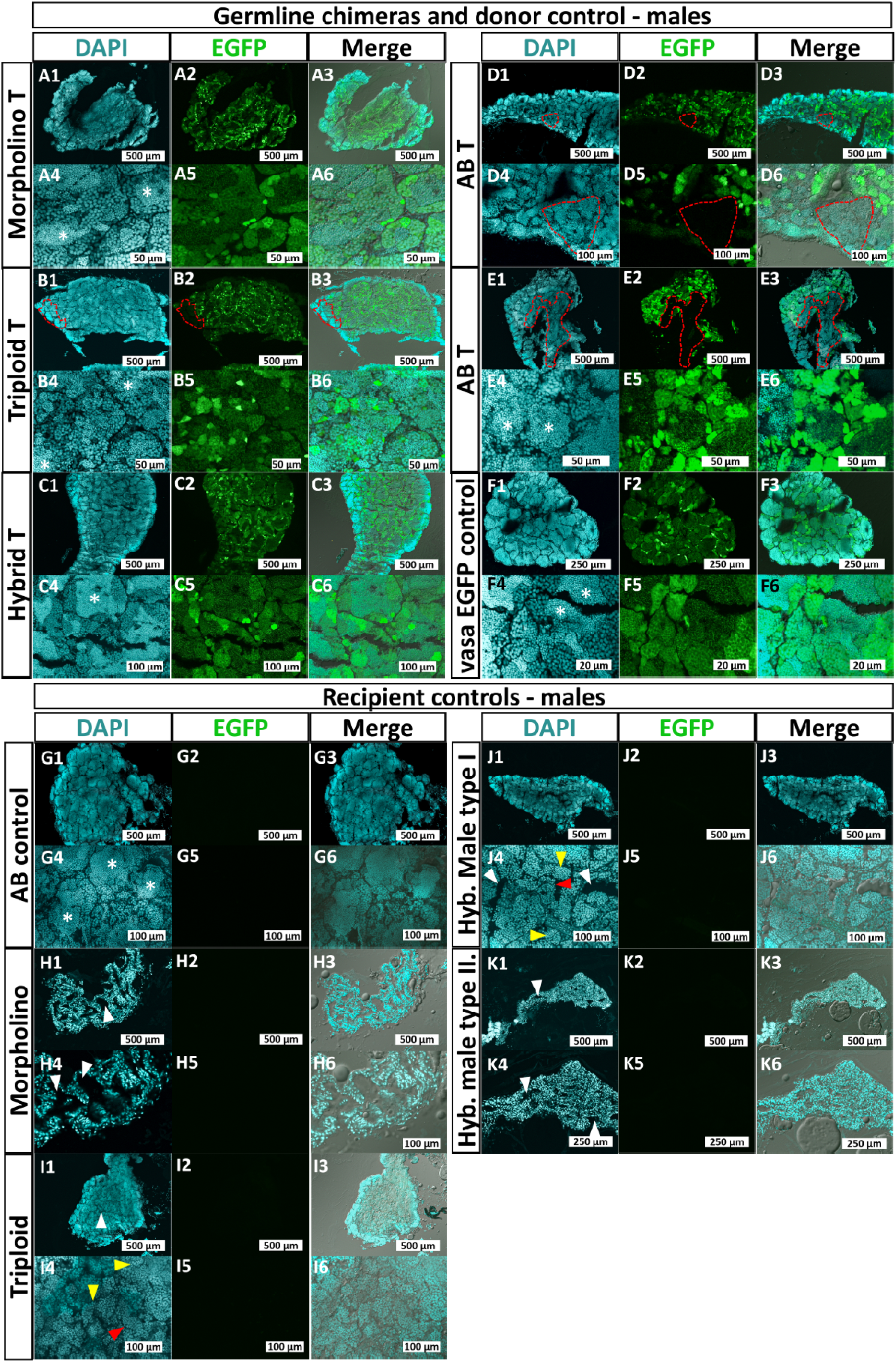
Distribution of transplanted germ cells from vas::EGFP donors in the recipients’ gonads. **A)** Morpholino treated male with exogenous spermatogenesis with EGFP signal occupying whole testis. **B)** Triploid male with most of the testes occupied by vas::EGFP positive spermatogenesis. B1-B3 red dashed line indicates spermatocysts with germ cells lacking EGFP signal. **C)** Hybrid male recipient with exogenous spermatogenesis occupying whole testis. **D, E)** AB transplanted males with only partial colonization of recipient testis by exogenous spermatogenesis. Note the red dashed lines depicting part of the testis with endogenous spermatogenesis lacking EGFP signal especially in E1-E3. **F)** vas::EGFP control specimen with well-organized spermatocysts and EGFP expression through whole section. **G)** AB control male with complete spermatogenesis and lumens filled with spermatozoa. **H)** Morpholino treated male lacking germ cells with developed empty lumens (white arrowheads). **I)** Triploid male specimen with developed testis with few individual spermatozoa (yellow arrowheads), poorly developed lumen (white arrowhead) and meiotic germ cells with aberrant morphology (red arrowhead). **J)** Hybrid male with developed testis and spermatogenesis. with few individual spermatozoa (yellow arrowheads), poorly developed lumen (white arrowhead) and meiotic germ cells with aberrant morphology (red arrowhead). **K)** Hybrid male with undeveloped testis lacking germ cells only with empty lumens (white arrowheads). White asterisks indicate lumen filled with spermatozoa. Note that EGFP signal intensity is strongest in the early-stage germ cells and decreasing by differentiation towards spermatozoa.

**Figure 8.**
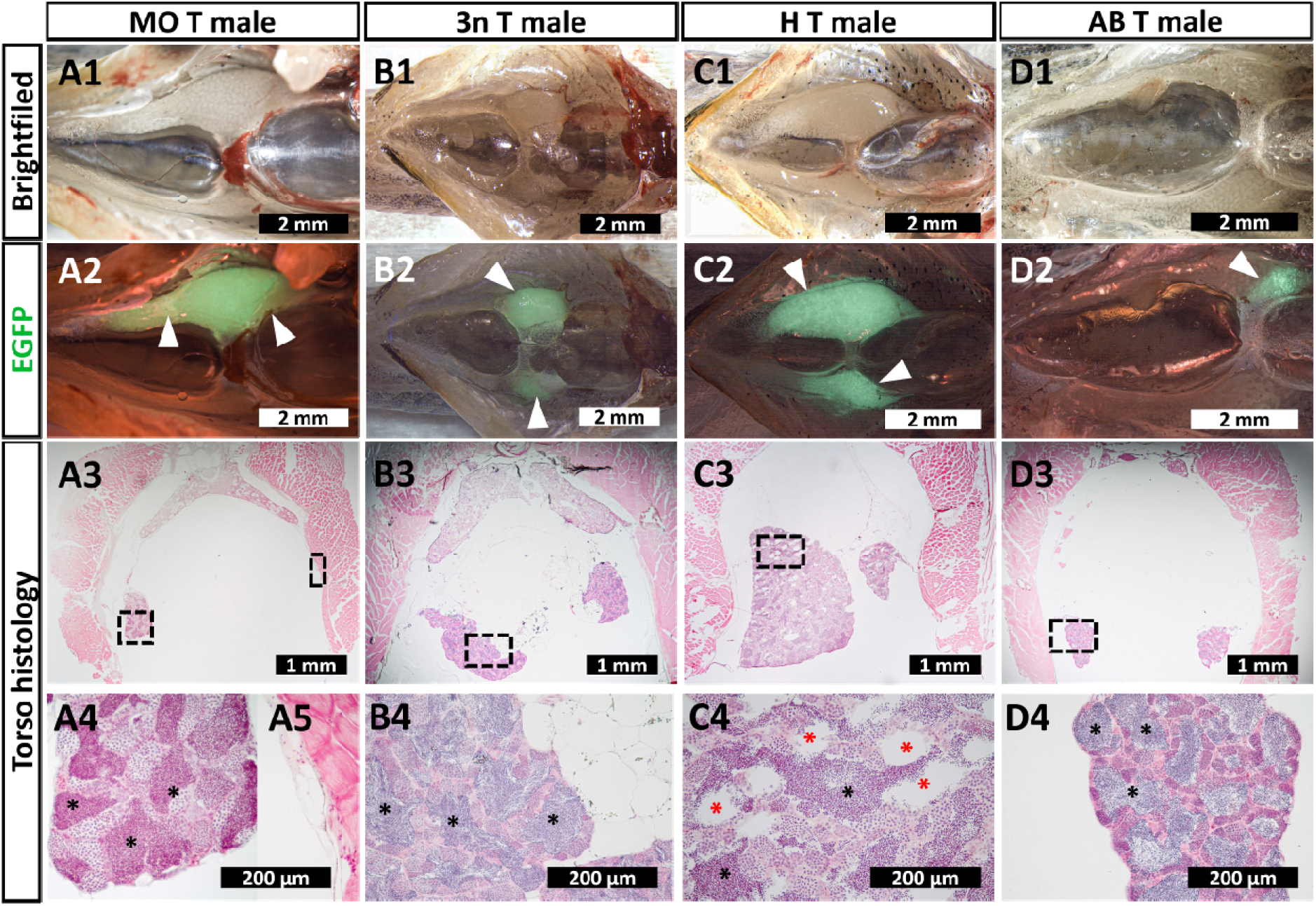
Gonadal development in germline chimeras. A) MO T chimeric male. A1-2) Colonized testis is apparent in the opened body cavity, while the second testis is very thin. Histological sections are showing colonized testis with filled lumens (black asterisks) (A4), while the second testis has typical structure of germ cell free gonad (A5). B) 3n T chimeric male. Both left and right testis are colonized in the medial/anterior part (B1-2). Histological sections are showing lumens filled with spermatozoa suggesting presence of spermatogenesis from donor-derived cells. C) H T chimeric male. C1-2) Developed testis are large and expressing EGFP signal. However, histological sections (C3-4) shows that many lumens are empty (red asterisk) suggesting that the encompassing germ cells are not undergoing proper gametogenesis D) AB T chimeric males with testis colonized in the very anterior part (D2). Testes are well filled with spermatozoa (D3-4).

### 3.4 Occurrence of chimeric females producing donor-derived eggs

Several females from AB T group were identified to have EGFP signal in their ovaries. Those fish were attempted for spawning with control AB males. Production of viable donor-derived eggs in zebrafish was confirmed in 7 from 10 spawned females. Individual AB T females produced EGFP positive eggs in various ratios, but their proportion was significantly lower compared to recipient-derived eggs (Fig. 9A). Eggs and later embryos from donor-derived EGFP eggs showed similar viability to the recipient-derived (endogenous) eggs. (Fig. 9B). The presence of oogenesis derived from transplanted male GSCs was also confirmed on ovarian cryosection by detection of EGFP signal (Fig. 10). Interestingly, overall sex ratio in AB T compared to AB C was slightly biased in favour of females (Fig. 9D) but without significant difference.

**Figure 9.**
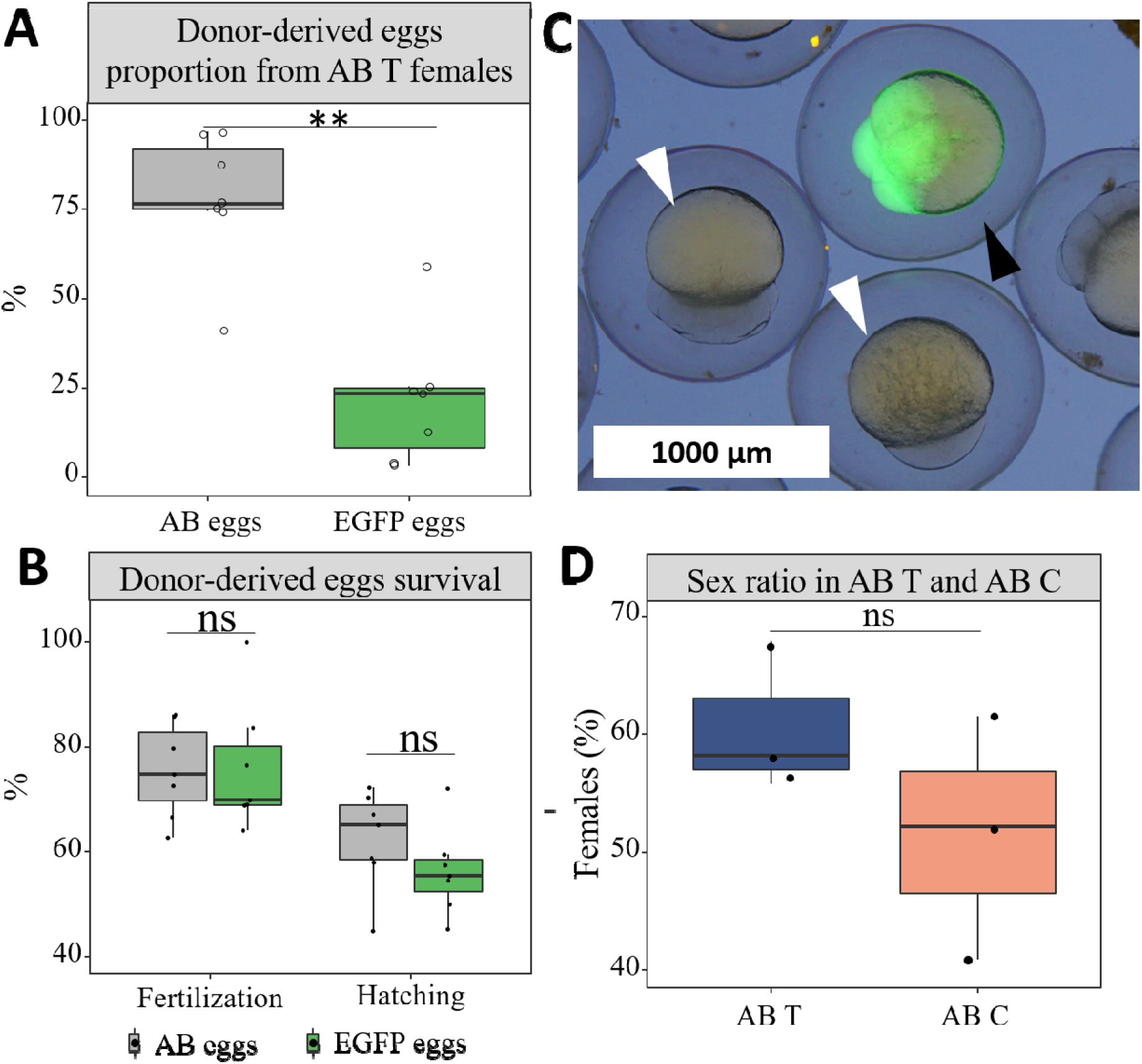
Female germline chimeras in AB T group. A) Proportion of produced recipient-derived and donor-derived oocytes. B) Survival rate of recipient-derived and donor-derived oocytes. C) Example of donor derived oocyte with strong EGFP signal (white arrowhead) and recipient-derived oocytes (black arrowhead) produced by AB T female. D) Overall incidence of females in AB T and AB C group. Asterisk stands for statistically significant difference (T-test, * * P < 0.01), while “ns” stand for no statistical difference (T-test, P > 0.05).

**Figure 10.**
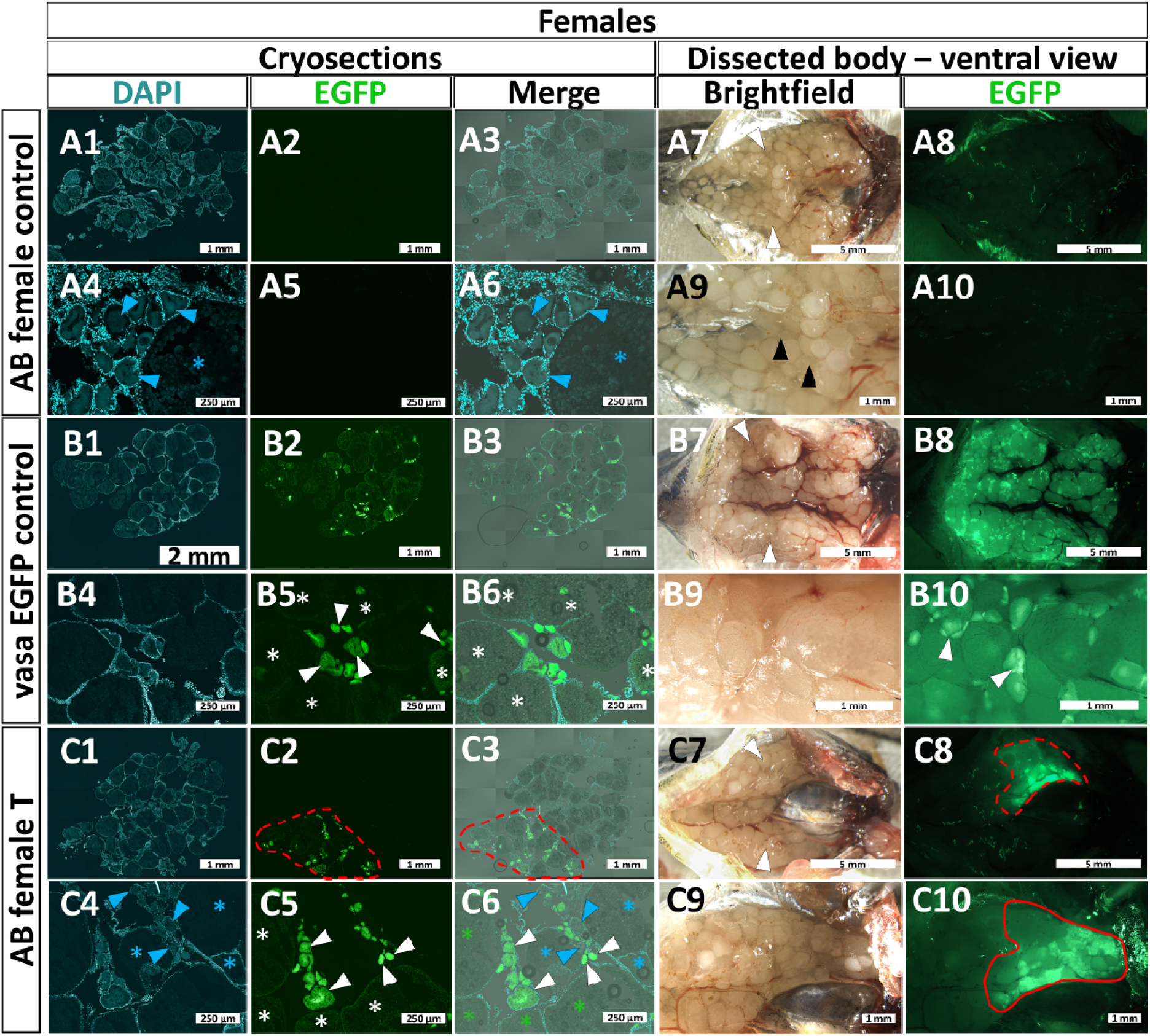
Detection of chimeric gonads in non-sterilized AB females. **A)** Control female from AB strain, only DAPI signal is detected on the whole ovary cryosections (A1-A6), no EGFP signal is detected in opened body cavity (A7-A10). **B)** Control female from vas::EGFP donor strain, DAPI as well EGFP signal is detected on the whole section of the ovaries. Early stage (small) oocytes have strong EGFP signal which is apparent on cryosections (B1-B6) as well on view on the opened body cavity (B7-B10). **C)** AB female germline chimera, ovarian germ cells derived from transplanted spermatogonia are occupying considerable part of the ovary indicated by red dashed line (C2-C3, C8 and C10). Magnified view on cryosection (C6) shows that endogenous oocytes positive only for DAPI signal (small oocytes indicated by blue arrow, advanced oocytes by blue asterisk) are developing in close contact with exogenous EGFP positive oocytes (small oocytes indicated by white arrow, advanced oocytes by white asterisk). View on opened body cavity shows anterior localization of donor derived oocytes indicated by red dashed line (C8 and C10). Images with 1,2 and 3 numerals were stitched from XY stack and images with 4, 5 and 6 numerals are magnified captions respectively. Images with 9 and 10 numerals are magnified captions of images with 7 and 8 numerals respectively.

### 3.5 Reproductive performance of chimeric males

The sperm concentration and total amount of produced sperm in germline chimeras was influenced by the fact that the testes comprised of donor-derived germ cells are not reaching their full size compared to controls. All sterilization methods interfered with the sperm motility, curvilinear and straight-line velocity (11A-C) which were usually significantly lower than in AB C group. This fact is clearly visible in Fig. 11D, where the largest proportion of fast spermatozoa were found in the recipient control. Only MO T group retained statistically comparable level of motility to donor strain and outperformed 3n and H T group, yet without significant differences. Also, it was apparent that motility performance in 3n T and H T groups was more dispersed showing very well and poorly performing males compared to other assessed groups. Results showed that MO recipients males produced highest volume of sperm (Fig. 11E), concentration of spermatozoa (Fig. 11F), total number of spermatozoa (Fig. 11G) and finally also total motile spermatozoa (Fig. 11H) among tested sterilized recipients. Overall results from sperm analysis are given in Figure 11.

**Figure 11.**
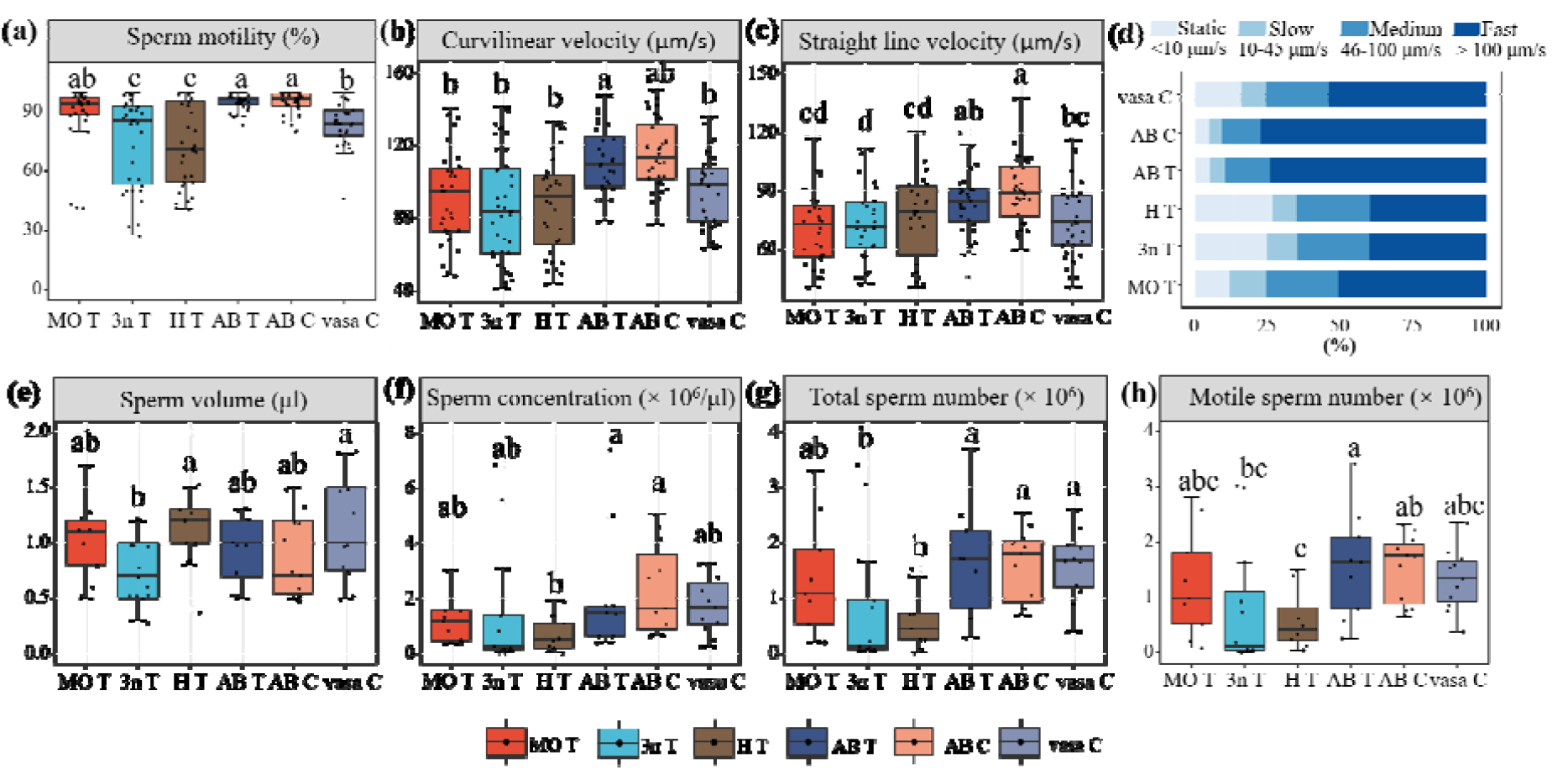
Reproductive performance of chimeric males with donor and recipient controls. Motility rate (%) (a), curvilinear velocity (μm/s) (b), straight-line velocity (μm/s) (c), percentage of sperm motility from total motility of spermatozoa evaluated at 15 s of PSA: rapid motility (>100 μm/s), medium motility (46 to 100 μm/s), slow motility (10 to 45 μm/s) and static spermatozoa (<10 μm/s) of tested groups at 15 s post activation. Total sperm volume collected from males (e), sperm concentration (× 10^6^/µl), total sperm number (× 10^6^) and total motile sperm number (× 10^6^). Values with a different lowercase letter are significantly different (P < 0.05, one-way analysis of variance (ANOVA) followed by an LSD test for post hoc multiple comparisons).

### 3.6 Fertilization trials

Semi artificial fertilization trials conducted individually (one experimental male with one control female) resulted in similar success of males to induce oviposition (number of spawning females) when over 70% of pairs attempted for spawning actually spawned (Fig. 12A), and number of oviposited eggs (Fig. 12B) was statistically comparable between transplanted groups and their respective controls (e.g. MO T and MO C group). Non-transplanted (sterile) MO C, 3n C and H C controls were able to induce oviposition, but no surviving progeny (reaching swim up stage) was obtained when most oviposited eggs were unfertilized or died during embryonic development. Comparison of transplanted recipients with AB and vas::EGFP controls showed poor performance of H T recipients, which was especially prominent in semi-artificial fertilization trials, while MO T and 3n T males showed performance comparable to one of the controls (AB C or vas::EGFP) (Fig. 12C). In vitro fertilization resulted in higher progeny production in all groups including controls (Fig. 12D). Importantly, the percentage of swim-up larvae was statistically comparable amongst all groups except the H T group.

**Figure 12.**
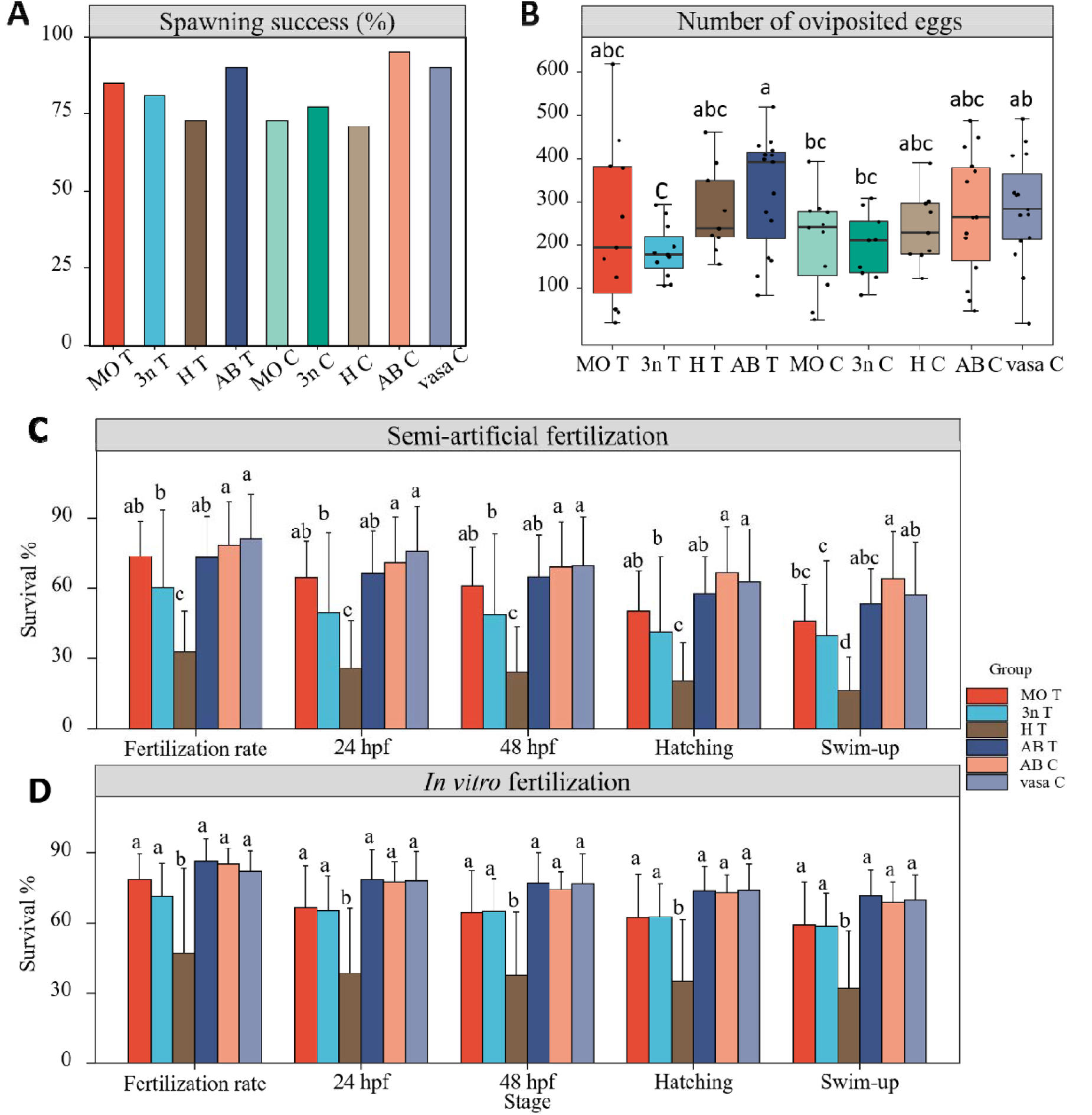
Reproductive success of chimeric males. A) Spawning success of males from experimental and control groups including sterile controls. Successful spawning of the given pair was recorded when 20 and more eggs were observed. B) Number of oviposited eggs during semi-artificial fertilization. Survival rates after semi-artificial (C) and in vitro fertilization tests (D) (mean ± S.D.). Values with different letters are significantly different among all groups (B) and within each development stage (C and D) (P < 0.05, one-way analysis of variance (ANOVA) followed by an LSD test for post hoc multiple comparisons).

Genotyping of hatched larva originating from crosses of chimeric males and AB control females showed 100% germline transmission detected by EGFP specific primers in MO T, 3n T and H T group. In the AB T group, only a low germline transmission rate was detected 12 ± 6 % (Mean ± SD). The complete dataset for individual males and their germline transmission rate is in Supplementary file 1.

## 4. DISCUSSION

The present study aimed to thoroughly assess different sterilization strategies for surrogacy in fish using zebrafish model, and their consequences on the reproductive output. We tested three types of sterilization for intraperitoneally transplanted zebrafish spermatogonia to help with the direct selection of the sterilization methods for species, which are not fully established in the laboratories. Similar kind of the study presented here (from recipient embryo to donor-derived gamete production) would be difficult to conduct in species such as tuna or sturgeons.

One of the first issues in choosing a convenient recipient is its availability and survival, which goes hand in hand with necessary efforts to achieve the given type of sterilization. Hybridization and triploidization are methods of choice for large scale sterility induction. However, further problems can appear in adult chimeras since their endogenous germ cells can proceed through gametogenesis. PGCs depletion during early embryonic development by targeting *dnd* gene requires precise injection of each embryo which is time consuming, demanding and suitable only for certain types of experiments because developing embryos have to be micromanipulated within limited time.

Each sterilization strategy has its pros and cons, which needs to be considered and evaluated carefully. Besides differently altered survival dependent upon the sterilization method, we were able to identify striking differences in gonadal development and reproductive output of differently sterilized surrogates. Once the gonads are free of endogenous GCs, it is likely to obtain the most consistent results without interference with gonadal development because there is no competition between exo- and endogenous GCs. Later observed reproductive differences might be the ultimate decisive factor for choosing the most perspective sterilization strategy to obtain germline chimeras with closest characteristics to the original donor strain.

The sterilization treatments negatively influenced survival prior to transplantation. Heat shock for triploidy induction and MO injection had a severe impact on the survival rate. This can be considered as expected based on previous results (Delomas and Dabrowski, 2018; Franěk et al., 2019b). Low survival due to the temperature treatment for triploidization group needs to be considered, however, it can be mitigated by using large amount of fertilized eggs. Situation with low survival in MO group is more challenging to be tackled since the number of injected embryos is limited by the skills of the personnel performing microinjection and developmental speed of fertilized eggs. However, it is also possible to alternate laborious MO delivery by microinjection using immersion of fertilized zebrafish eggs *in vivo* MO (capable of penetration and transport through cell membranes) (Wong and Zohar, 2015).

Post-transplantation survival is also an important aspect for the selection of suitable recipients and sterilization strategy. MO T and 3n T groups with their respective controls showed comparable post-transplantation survival to control groups from donor and recipient strains over the duration of the experiment. Usually, larvae malformed due to the sterilization treatments could not proceed embryogenesis or they did not reach swim-up feeding stage. Thus, only healthy and feeding larvae are used for transplantation and they do not further interfere with survival in case of MO and 3n recipients. H T group experienced the lowest survival post transplantation. We presume that this can be attributed to the hybridization itself caused by partial genetic incompatibility of parental species. Also, influence of transplanted cells is possible, since H C group showed higher post-transplantation survival. Interestingly, comparison between transplanted and control group in MO and 3n recipients showed that transplantation procedure did not interfere with the survival rate. Lower robustness of the ZFxPD hybrids was probably challenged by transplantation. In general, post-transplantional survival should be of larger concern than survival before transplantation. Due to low survival in the H T group, some germline chimeras were lost during ongrowing. This can represent a severe issue when amount of available donor’s GSCs is limited. In other words, it is more reasonable to sacrifice lower survival during embryogenesis for the sake of the post-transplantation survival until adulthood.

All tested recipients developed gonads capable to support transplanted GSCs including non-sterilized AB recipients. Transplantation success evaluated by the colonization rate was found as only a preliminary indicator because it did not show differences among assessed groups, which were later obvious in adult fish. Although we have used sterilization treatment including complete PGCs ablation, transplantation into non-sterilized AB recipients still resulted in the EGFP positive cells in more than 30% of the transplanted fish (2wpt). This finding clearly shows that colonization rate assessed few days or weeks post-transplantation does not guarantee high success since incidence of chimeric gonads in non-sterile AB T adults was indeed low. Similar pattern was also observed in hybrid recipients showing high colonization rate but low incidence of adult germline chimera. However, the low number of adult germline chimera in H T was attributed to their low survival during the experiment. In other species it was also documented that recipient’s gonads with endogenous GCs can be colonized with transplanted cells. However, introduced cells are later losing the pace of the recipient’s gametogenesis and are finally outcompeted (Yoshizaki et al., 2016). There are probably two scenarios for transplanted GSCs, which are dependent on the sterility level and determine the success of the transplanted cells. First, when the GSCs are introduced into the PGCs depleted gonads, they are not in competition for the space (germinal niches) and transplantation success can be evaluated early. The second one takes place in PGCs non-depleted recipients (non-sterile control, 3n or H) when introduced cells colonized the gonad, however, introduced GSCs might be limited until endogenous GCs proceed to affected gametogenesis stage. If endogenous GCs do not experience developmental problems (e.g. arrest in meiosis), the relative proportion of exogenous cells decreases, and they cannot further occupy more testicular niches. We presume that eventual loss of transplanted cells in the competitive environment of the unsterile gonads takes place during more advanced stages of the gonadal development. It would be very informative to identify this period precisely and to find mechanistic and molecular reasons behind this loss of the introduced cells and later use it for interventions to increase transplantation success.

Importantly, PGCs depletion and triploidization treatment clearly showed to promote chimeric gonad incidence and development in adults. This finding is striking, especially in triploids since they have quite well-developed testes with complete, although impaired, spermatogenesis with lower GCs numbers. Thus, we suggest that fish GSCs are in the host environment opportunistic and capable of utilizing the gonadal environment once they experience some developmental problems in gametogenesis. Therefore, the extent of gonadal development in the sterile host might not be always crucial for surrogacy success. Incidence of adult AB T male chimeras was about 10% and transplanted cells were capable of establishing spermatogenesis in tiny part of the testes resulting in low production of donor-derived sperm reflected by low germline transmission rate. On the other hand, dissection of all sterilized recipients showed potential of the transplanted cells to establish and expand spermatogenesis on a large scale. Macroscopically, MO T group showed the most developed chimeric testes. 3n T and H T group could also create a considerable area with EGFP positive cells. However, full bilateral or unilateral development of chimeric testes was rare in 3n T an H T groups, when cells were localized rather spatially not occupying full length of the testes. These findings lead us to presumption about competition between exo- and endogenous germ cells, which has further consequences to the proportion of adult germline chimeras and their reproductive performance.

Reasons causing low sperm performance in 3n T and H T groups are challenging to interpret. Presence of recipient-derived sperm in HT was confirmed in the testicular lumen by histology; however we roughly estimated less than 1% incidence of abnormally sized spermatozoa. At light and fluorescent microscopy level, we could identify only few individual abnormally sized and EGFP negative spermatozoa among hundreds of spermatozoa showing donor-derived characteristics. Therefore, it is not probable, that overall low sperm performance in HT group was caused by recipient-derived sperm. Partial genetic incompatibility of parental species has clear consequences on endogenous gametogenesis. Although transplanted cells could establish normal spermatogenesis, some molecular alteration conditioning final spermatogenic stages might be present, and in turn, resulted in poor performance of some 3n T and H T chimeric males. Hybrid testes likely need to cope with increased apoptosis of germ cells, as shown previously due to failure in homologous chromosome pairing (Ponjarat et al., 2019). We can speculate that pathways responsible for removing defective germ cells in the hybrid and triploid testes have further negative consequences on the final spermiogenesis phase, causing low sperm performance in some males.

Observed differences in sperm quality and quantity were reflected in both fertilization trials. Importantly, lower sperm performance and fertilization success in semi-artificial tests was less striking during *in vivo* fertilization trials. The potential problem and risks of 3n and especially H recipients are dispersed motility rates. About 25% of spermatozoa from 3n T and H T male chimeras had extremely low velocity (<10 μm/s). Moreover, presence of well performing males as well as bad performing males was evident. Sperm performance in 3n T and H T groups in the combination of sperm quantity and motility indicates the fact that these groups produce on average about 5 to 20 times less motile spermatozoa than the MO T group. Sperm from low performing males can compromise well-performing sperm once they are pooled during collection. It is important for *in vitro* fertilization, always collecting sperm individually and pooling it only when fertilizing the eggs. If the sperm is pooled immediately after sperm collection, it rapidly decreases sperm motility and fertility (Cheng et al. 2021).

To our surprise, females from AB T group produced donor-derived eggs giving rise to viable embryos. This finding represents first report on surrogate egg production in zebrafish, because triploid or PGCs depleted zebrafish are all male only (Delomas and Dabrowski, 2018; Slanchev et al., 2005). Proportion of produced donor-derived EGFP positive eggs was rather low (about 20% on average). Therefore, it is clear that non-sterilized ovaries constitute very competitive environment. Interestingly, total proportion of chimeric males and chimeric females in AB T group was 19 and 17 individuals, respectively. This finding suggests that the trans-differentiation of male GSCs to female GSCs in the ovarian environment is not decisive for successful intraspecific surrogacy.

Interestingly, incidence of hybrid females in control groups was rare and no chimeric hybrid female was observed in this study. Similarly, Wong and Saito (2011) also did not record any hybrid chimeric females after ovarian cell transplantation. In overall, low incidence of hybrid females in danio species was recently reported in cross of zebrafish and spotted danio (*Danio nigrofasciatus*) (Endoh et al., 2020). It is evident that female hybrids are likely to experience more severe gametogenesis alteration than males. Therefore, it is reasonable to expect that poorly developed ovaries in hybrid females cannot provide proper environment for transplanted cells and production of donor-derived zebrafish eggs can be achieved only through non-sterilized female recipients as it is for the first time described in the present study.

Predictable and stable gonadal phenotype development was identified as a concern in hybrid male recipients when three distinct phenotypes were observed. Previous study utilizing ZF x PD hybrid recipients showed lack of spermatozoa in the hybrid testes (Wong and Saito, 2011) while our study confirmed that hybrid GCs are capable to proceed throughout entire spermatogenesis resulting in the production of extremely abnormal spermatozoa. On the other hand, gonadal phenotypes in PGC ablated fish and triploids were consistent. Fish produced by *dnd* gene targeting developed empty gonads composed of the solely somatic cells. Further female or male fate differentiation of the sterile gonad is species specific. Germ cell less zebrafish and medaka have been shown to develop into phenotypical males only (Kurokawa et al., 2007; Slanchev et al., 2005; Tzung et al., 2015). Otherwise, several species have germ cell independent sex differentiation such as loach (Fujimoto et al., 2010), goldfish (Goto et al., 2012), trout (Yoshizaki et al., 2016), Atlantic salmon (Wargelius et al., 2016) or rosy bitterling (Octavera and Yoshizaki, 2018). Zebrafish (Franěk et al., 2019) and rosy bitterling surrogates (Octavera and Yoshizaki, 2018) showed only male development after germ cell transplantation, meaning that introduced additional GSCs are not capable of rescuing female fate of the gonad. Majority of induced triploid surrogates can differentiate into both sexes, including salmonids (Lee et al., 2013; Okutsu et al., 2007), medaka (Seki et al., 2017), grass puffer (*Takifugu niphobles*) (Hamasaki et al., 2017), or Nibe croaker (*Nibea mitsukurii*) (Yoshikawa et al., 2017). Hybrids in this study showed gonadal phenotypes with developed testes, altered ovaries, and empty gonads resembling germ cell-depleted MO phenotype.

Strain specific differences or different age of the assessed fish can probably play an important role in gonadal phenotype of hybrids. Similar variance in testicular phenotype was described in mackerel hybrid of *Scomber australasicus* x *S. japonicus*, when part of the hybrid males could proceed through spermatogenesis while the second phenotype was germ cell less (Kawamura et al., 2020). Consequently, semi fertility of the hybrid could interfere with the fertilization since endo- and exogenous gametes are in the competition for the ova. Therefore, suitability of hybrids for surrogacy needs to be verified thoroughly in particular species. The hybridization itself is very convenient tool for recipient production for surrogacy because it requires only fertilization without further manipulation. However, since mechanisms causing occurrence of sterile and GCs producing gonadal phenotypes are unknown it should be evaluated with cautions.

## CONCLUSION

GSCs manipulation is potent biotechnology to ameliorate breeding of aquaculture species and preserve valuable genetic resources in environmentally relevant or even endangered species. This study aimed to identify best sterilization treatment - essential factor influencing the surrogacy success rate. The presented study assessed various sterilization treatments in fish for surrogates preparation and their influence on gonadal development and reproductive output in germline chimeras. Of the utmost importance, germ cell-free gonads were identified as the best environment for transplanted cells yielding the highest transplantation success and gonadal development. Importantly, reproductive performance of males including quantity and motility parameters and fertilization rate clearly favors germ cell depleted recipients. The use of triploid and hybrid males from the point of view of the production of sufficient quantity and quality sperm proves to be risky to achieve stable results. Moreover, only germ cell depleted recipient retained reproductive characteristics of the donor strain. Presented findings should help in decision on what type of sterilization should be used prior to transplantation and surrogacy induction, especially in non-model fish species.

The overall suitability and versatility of the zebrafish surrogate model can be utilized to provide deeper insights into the mechanism of GCs behaviour in the recipient’s gonads and dissect specific factors influencing promotion of the exogenous GCs development. Our interest should also be directed to the molecular aspects of surrogacy. Nowadays, GSCs manipulations and surrogacy were performed in wide range of species. However, we know only little about the lifetime or transgenerational consequences of gametes produced from surrogate parents and how they can possibly influence resulting progeny and its performance.

## FUNDING

The work was supported by National Agriculture Agency project number QK1910428, by the Ministry of Education, Youth and Sports of the Czech Republic - project Biodiversity (CZ.02.1.01/0.0/0.0/16_025/0007370). This project has received funding from the European Union’s Horizon 2020 Research and Innovation Programme under grant agreement No 652831 (AQUAEXCEL2020) and No 871108 (AQUAEXCEL3.0). This output reflects only the authors’ view and the European Union cannot be held responsible for any use that may be made of the information contained therein.

## AUTHOR CONTRIBUTIONS

Conceptualization: RF; Data curation and Formal Analysis: YC; Funding acquisition: VK, OL, IŠ, MP; Investigation: RF, YC, MF, XX, MAS, VK, OL, IŠ, MP; Methodology: RF, YC, IŠ; Project administration: RF; Resources: RF, YC, MF, OL, IŠ, MP; Supervision: RF; Validation: RF, YC; Visualization: RF, YC; Writing – original draft: RF; Writing – review & editing: all authors contributed.

## Notes

### Competing Interest Statement

The authors have declared no competing interest.

## REFERENCES

Baloch, A.R., Franěk, R., Saito, T., Martin, Pšenička., 2019a. Dead-end (dnd) protein in fish — a review. Fish Physiol. Biochem. 47, 777–784. https://doi.org/10.1007/s10695-018-0606-x

Baloch, A.R., Franěk, R., Tichopád, T., Fučíková, M., Rodina, M., Pšenička, M., 2019b. Dnd1 Knockout in Sturgeons By CRISPR/Cas9 Generates Germ Cell Free Host for Surrogate Production. Animals 9, 174. https://doi.org/10.3390/ani9040174

Cheng, Y., Franěk, R., Rodina, M., Xin, M., Cosson, J., Zhang, S., Linhart, O., 2021. Optimization of Sperm Management and Fertilization in Zebrafish (Danio rerio (Hamilton)). Animals 11, 1558. https://doi.org/10.3390/ANI11061558

Delomas, T.A., Dabrowski, K., 2018. Why are triploid zebrafish all male? Mol. Reprod. Dev. 85, 612–621. https://doi.org/10.1002/mrd.22998

Endoh, M., Shima, F., Havelka, M., Asanuma, R., Yamaha, E., Fujimoto, T., Arai, K., 2020. Hybrid between Danio rerio female and Danio nigrofasciatus male produces aneuploid sperm with limited fertilization capacity. PLoS ONE 15, e0233885. https://doi.org/10.1371/journal.pone.0233885

Franěk, R., Kašpar, V., Shah, M.A., Gela, D., Pšenička, M., 2021. Production of common carp donor-derived offspring from goldfish surrogate broodstock. Aquaculture 534, 736252. https://doi.org/10.1016/j.aquaculture.2020.736252

Franěk, R, Marinović, Z., Lujić, J., Urbányi, B., Fučíková, M., Kašpar, V., Pšenička, M., Horváth, Á., 2019a. Cryopreservation and transplantation of common carp spermatogonia. PLOS ONE 14, e0205481. https://doi.org/10.1371/journal.pone.0205481

Franěk, R., Tichopád, T., Steinbach, C., Xie, X., Lujić, J., Marinović, Z., Horváth, Á., Kašpar, V., Pšenička, M., Lujić, J., Horváth, Á., Pšenička, M., 2019b. Preservation of female genetic resources of common carp through oogonial stem cell manipulation. Cryobiology 87, 78–85. https://doi.org/10.1016/j.cryobiol.2019.01.016

Franěk, R., Tichopád, T., Fučíková, M., Steinbach, C., Pšenička, M., 2019c. Production and use of triploid zebrafish for surrogate reproduction. Theriogenology 140, 33–43. https://doi.org/10.1016/J.THERIOGENOLOGY.2019.08.016

Fujimoto, T., Nishimura, T., Goto-Kazeto, R., Kawakami, Y., Yamaha, E., Arai, K., 2010. Sexual dimorphism of gonadal structure and gene expression in germ cell-deficient loach, a teleost fish. Proc. Natl. Acad. Sci. U. S. A. 107, 17211–17216. https://doi.org/10.1073/pnas.1007032107

Fujimoto, T., Yasui, G.S., Yoshikawa, H., Yamaha, E., Arai, K., 2008. Genetic and reproductive potential of spermatozoa of diploid and triploid males obtained from interspecific hybridization of Misgurnus anguillicaudatus female with M. mizolepis male. J. Appl. Ichthyol. 24, 430–437. https://doi.org/10.1111/j.1439-0426.2008.01131.x

Goto, R., Saito, T., 2019. A state-of-the-art review of surrogate propagation in fish. Theriogenology 133, 216–227. https://doi.org/10.1016/j.theriogenology.2019.03.032

Goto, R., Saito, T., Takeda, T., Fujimoto, T., Takagi, M., Arai, K., Yamaha, E., 2012. Germ cells are not the primary factor for sexual fate determination in goldfish. Dev. Biol. 370, 98–109. https://doi.org/10.1016/j.ydbio.2012.07.010

Hamasaki, M., Takeuchi, Y., Yazawa, R., Yoshikawa, S., Kadomura, K., Yamada, T., Miyaki, K., Kikuchi, K., Yoshizaki, G., 2017. Production of Tiger Puffer Takifugu rubripes Offspring from Triploid Grass Puffer Takifugu niphobles Parents. Mar. Biotechnol. 19, 579–591. https://doi.org/10.1007/s10126-017-9777-1

Hattori, R.S., Yoshinaga, T.T., Katayama, N., Hattori-Ihara, S., Tsukamoto, R.Y., Takahashi, N.S., Tabata, Y.A., 2019. Surrogate production of Salmo salar oocytes and sperm in triploid Oncorhynchus mykiss by germ cell transplantation technology. Aquaculture 506, 238–245. https://doi.org/10.1016/J.AQUACULTURE.2019.03.037

Iwasaki-Takahashi, Y., Shikina, S., Watanabe, M., Banba, A., Yagisawa, M., Takahashi, K., Fujihara, R., Okabe, T., Valdez, D.M., Yamauchi, A., Yoshizaki, G., 2020. Production of functional eggs and sperm from in vitro-expanded type A spermatogonia in rainbow trout. Commun. Biol.. 3, 308. https://doi.org/10.1038/s42003-020-1025-y

Jin, Y.H., Robledo, D., Hickey, J., McGrew, M., Houston, R., 2021. Surrogate broodstock to enhance biotechnology research and applications in aquaculture. Biotechnol. Adv. 49, 107756. https://doi.org/10.1016/j.biotechadv.2021.107756

Kawakami, Y., Goto-Kazeto, R., Saito, T., Fujimoto, T., Higaki, S., Takahashi, Y., Arai, K., Yamaha, E., 2010. Generation of germline chimera zebrafish using primordial germ cells isolated from cultured blastomeres and cryopreserved embryoids. Int. J. Dev. Biol. 54, 1491–1499. https://doi.org/10.1387/ijdb.093059yk

Kawamura, W., Tani, R., Yahagi, H., Kamio, S., Morita, T., Takeuchi, Y., Yazawa, R., Yoshizaki, G., 2020. Suitability of hybrid mackerel (Scomber australasicus × S. japonicus) with germ cell-less sterile gonads as a recipient for transplantation of bluefin tuna germ cells. Gen. Comp. Endocrinol. 295, 113525. https://doi.org/10.1016/j.ygcen.2020.113525

Kurokawa, H., Saito, D., Nakamura, S., Katoh-Fukui, Y., Ohta, K., Baba, T., Morohashi, K.I., Tanaka, M., 2007. Germ cells are essential for sexual dimorphism in the medaka gonad. Proc. Natl. Acad. Sci. U. S. A. 104, 16958–16963. https://doi.org/10.1073/pnas.0609932104

Lacerda, S.M.S.N., Batlouni, S.R., Costa, G.M.J., Segatelli, T.M., Quirino, B.R., Queiroz, B.M., Kalapothakis, E., França, L.R., 2010. A new and fast technique to generate offspring after germ cells transplantation in adult fish: The nile tilapia (Oreochromis niloticus) model. PLoS ONE 5, e10740. https://doi.org/10.1371/journal.pone.0010740

Lee, S., Iwasaki, Y., Shikina, S., Yoshizaki, G., 2013. Generation of functional eggs and sperm from cryopreserved whole testes. Proc. Natl. Acad. Sci. U. S. A. 110, 1640–1645. https://doi.org/10.1073/pnas.1218468110

Li, Q., Fujii, W., Naito, K., Yoshizaki, G., 2017. Application of dead end-knockout zebrafish as recipients of germ cell transplantation. Mol. Reprod. Dev. 84, 1100–1111 https://doi.org/10.1002/mrd.22870

Linhartová, Z., Saito, T., Kašpar, V., Rodina, M., Prášková, E., Hagihara, S., Pšenička, M., 2015. Sterilization of sterlet Acipenser ruthenus by using knockdown agent, antisense morpholino oligonucleotide, against dead end gene. Theriogenology 84, 1246–1255. https://doi.org/10.1016/j.theriogenology.2015.07.003

Marinović, Z., Li, Q., Lujić, J., Iwasaki, Y., Csenki, Z., Urbányi, B., Yoshizaki, G., Horváth, Á., 2019. Preservation of zebrafish genetic resources through testis cryopreservation and spermatogonia transplantation. Sci. Rep. 9, 13861. https://doi.org/10.1038/s41598-019-50169-1

Murray, D.S., Kainz, M.J., Hebberecht, L., Sales, K.R., Hindar, K., Gage, M.J.G., 2018. Comparisons of reproductive function and fatty acid fillet quality between triploid and diploid farm Atlantic salmon (Salmo salar). R. Soc. Open Sci. 5, 180493. https://doi.org/10.1098/RSOS.180493

Nagasawa, K., Ishida, M., Octavera, A., Kusano, K., Kezuka, F., Kitano, T., Yoshiura, Y., Yoshizaki, G., 2019. Novel method for mass producing genetically sterile fish from surrogate broodstock via spermatogonial transplantation. Biol. Reprod. 100, 535–546. https://doi.org/10.1093/biolre/ioy204

Nóbrega, R.H., Greebe, C.D., van de Kant, H., Bogerd, J., de França, L.R., Schulz, R.W., 2010. Spermatogonial stem cell niche and spermatogonial stem cell transplantation in zebrafish. PLoS ONE 5, e12808. https://doi.org/10.1371/journal.pone.0012808

Octavera, A., Yoshizaki, G., 2018. Production of donor-derived offspring by allogeneic transplantation of spermatogonia in Chinese rosy bitterling. Biol. Reprod. 100, 1108–1117. https://doi.org/10.1093/biolre/ioy236

Okutsu, T., Shikina, S., Kanno, M., Takeuchi, Y., Yoshizaki, G., 2007. Production of Trout Offspring from Triploid Salmon Parents. Science 317, 15–17.

Orr, H.A., Irving, S., 2001. Complex epistasis and the genetic basis of hybrid sterility in the Drosophila pseudoobscura Bogota-USA hybridization. Genetics 158, 1089–1100. https://doi.org/10.1093/genetics/158.3.1089

Piferrer, F., Beaumont, A., Falguière, J.C., Flajšhans, M., Haffray, P., Colombo, L., 2009. Polyploid fish and shellfish: Production, biology and applications to aquaculture for performance improvement and genetic containment. Aquaculture 293, 125–156. https://doi.org/10.1016/j.aquaculture.2009.04.036

Piva, L.H., de Siqueira-Silva, D.H., Goes, C.A.G., Fujimoto, T., Saito, T., Dragone, L.V., Senhorini, J.A., Porto-Foresti, F., Ferraz, J.B.S., Yasui, G.S., 2018. Triploid or hybrid tetra: Which is the ideal sterile host for surrogate technology? Theriogenology 108, 239–244. https://doi.org/10.1016/j.theriogenology.2017.12.013

Ponjarat, J., Singchat, W., Monkheang, P., Suntronpong, A., Tawichasri, P., Sillapaprayoon, S., Ogawa, S., Muangmai, N., Baicharoen, S., Peyachoknagul, S., Parhar, I., Na-Nakorn, U., Srikulnath, K., 2019. Evidence of dramatic sterility in F1 male hybrid catfish [male Clarias gariepinus (Burchell, 1822)=×=female C. macrocephalus (Günther, 1864)] resulting from the failure of homologous chromosome pairing in meiosis I. Aquaculture 505, 84–91. https://doi.org/10.1016/J.AQUACULTURE.2019.02.035

Ryu, J.H., Gong, S.P., 2020. Enhanced enrichment of medaka ovarian germline stem cells by a combination of density gradient centrifugation and differential plating. Biomolecules 10,1477. https://doi.org/10.3390/biom10111477

Saito, T., Goto-Kazeto, R., Fujimoto, T., Kawakami, Y., Arai, K., Yamaha, E., 2010. Inter-species transplantation and migration of primordial germ cells in cyprinid fish. Int. J. Dev. Biol. 54, 1479–1484. https://doi.org/10.1387/ijdb.103111ts

Seki, S., Kusano, K., Lee, S., Iwasaki, Y., Yagisawa, M., Ishida, M., Hiratsuka, T., Sasado, T., Naruse, K., Yoshizaki, G., 2017. Production of the medaka derived from vitrified whole testes by germ cell transplantation. Sci. Rep. 7, 43185. https://doi.org/10.1038/srep43185

Shikina, S., Nagasawa, K., Hayashi, M., Furuya, M., Yoshizaki, G., 2013. Short-term in vitro culturing improves transplantability of type A spermatogonia in rainbow trout (Oncorhynchus mykiss). Mol. Reprod. Dev. 80, 763–773, https://doi.org/10.1002/mrd.22208

Škugor, A., Tveiten, H., Krasnov, A., Andersen, Ø., 2014. Knockdown of the germ cell factor Dead end induces multiple transcriptional changes in Atlantic cod (Gadus morhua) hatchlings. Anim. Reprod. 144, 129–137. https://doi.org/10.1016/j.anireprosci.2013.12.010

Slanchev, K., Stebler, J., de la Cueva-Méndez, G., Raz, E., 2005. Development without germ cells: the role of the germ line in zebrafish sex differentiation. Proc. Natl. Acad. Sci. U. S. A. 102, 4074–4079. https://doi.org/10.1073/pnas.0407475102

Sullivan-Brown, J., Bisher, M.E., Burdine, R.D., 2011. Embedding, serial sectioning and staining of zebrafish embryos using JB-4 resin. Nat. Prot. 6, 46–55. https://doi.org/10.1038/nprot.2010.165

Takeuchi, Y., Yoshizaki, G., Takeuchi, T., 2003. Generation of Live Fry from Intraperitoneally Transplanted Primordial Germ Cells in Rainbow Trout. Biol. Reprod. 69, 1142–1149. https://doi.org/10.1095/biolreprod.103.017624

Tichopád, T., Vetešník, L., Šimková, A., Rodina, M., Franěk, R., Pšenička, M., 2020. Spermatozoa morphology and reproductive potential in F1 hybrids of common carp (Cyprinus carpio) and gibel carp (Carassius gibelio). Aquaculture 521, 735092. https://doi.org/10.1016/j.aquaculture.2020.735092

Tzung, K.W., Goto, R., Saju, J.M., Sreenivasan, R., Saito, T., Arai, K., Yamaha, E., Hossain, M.S., Calvert, M.E.K., Orbán, L., 2015. Early depletion of primordial germ cells in zebrafish promotes testis formation. Stem Cell Rep. 4, 61–73. https://doi.org/10.1016/j.stemcr.2014.10.011

Wargelius, A., Leininger, S., Skaftnesmo, K.O., Kleppe, L., Andersson, E., Taranger, G.L., Schulz, R.W., Edvardsen, R.B., 2016. Dnd knockout ablates germ cells and demonstrates germ cell independent sex differentiation in Atlantic salmon. Sci. Rep. 6, 21284. https://doi.org/10.1038/srep21284

Wong, T.-T., Saito, T., Crodian, J., Collodi, P., 2011. Zebrafish germline chimeras produced by transplantation of ovarian germ cells into sterile host larvae. Biol. Reprod. 84, 1190–1197. https://doi.org/10.1095/biolreprod.110.088427

Wong, T.-T., Zohar, Y., 2015. Production of reproductively sterile fish by a non-transgenic gene silencing technology. Sci. Rep. 5, 15822. https://doi.org/10.1016/j.ygcen.2014.12.012

Wong, T.T., Collodi, P., 2013. Inducible Sterilization of Zebrafish by Disruption of Primordial Germ Cell Migration. PLoS ONE 8, e68455. https://doi.org/10.1371/journal.pone.0068455

Xie, X., Li, P., Pšenička, M., Ye, H., Steinbach, C., Li, C., Wei, Q., 2019. Optimization of in vitro culture conditions of sturgeon germ cells for purpose of surrogate production. Animals 9, 106. https://doi.org/10.3390/ani9030106

Yang, Z., Yu, Y., Tay, Y.X., Yue, G.H., 2021. Genome editing and its applications in genetic improvement in aquaculture. Rev. Aquac.. https://doi.org/10.1111/RAQ.12591

Yoshikawa, H., Takeuchi, Y., Ino, Y., Wang, J., Iwata, G., Kabeya, N., Yazawa, R., Yoshizaki, G., 2017. Efficient production of donor-derived gametes from triploid recipients following intra-peritoneal germ cell transplantation into a marine teleost, Nibe croaker (Nibea mitsukurii). Aquaculture 478, 35–47. https://doi.org/10.1016/J.AQUACULTURE.2016.05.011

Yoshikawa, H., Xu, D., Ino, Y., Yoshino, T., Hayashida, T., Wang, J., Yazawa, R., Yoshizaki, G., Takeuchi, Y., 2018. hybrid sterility in fish caused by mitotic arrest of primordial germ cells. Genetics 209, 507–521. https://doi.org/10.1534/genetics.118.300777

Yoshizaki, G., Lee, S., 2018. Production of live fish derived from frozen germ cells via germ cell transplantation. Stem Cell Res. 29, 103–110. https://doi.org/10.1016/J.SCR.2018.03.015

Yoshizaki, G., Takashiba, K., Shimamori, S., Fujinuma, K., Shikina, S., Okutsu, T., Kume, S., Hayashi, M., 2016. Production of germ cell-deficient salmonids by dead end gene knockdown, and their use as recipients for germ cell transplantation. Mol. Reprod. Dev. 83, 298–311. https://doi.org/10.1002/mrd.22625

Yoshizaki, G., Yazawa, R., 2019. Application of surrogate broodstock technology in aquaculture. Fish. Sci. 85, 429–437. https://doi.org/10.1007/s12562-019-01299-y

Zhang, F., Li, X., Hao, Y., Li, Y., Ye, D., He, M., Wang, H., Zhu, Z., Sun, Y., 2021. Surrogate production of genome edited sperm from a different subfamily by spermatogonial stem cell transplantation. bioRxiv 2021.04.20.440715. https://doi.org/10.1101/2021.04.20.440715

Zhang, F., Li, X., He, M., Ye, D., Xiong, F., Amin, G., Zhu, Z., Sun, Y., 2020. Efficient generation of zebrafish maternal-zygotic mutants through transplantation of ectopically induced and Cas9/gRNA targeted primordial germ cells. J. Genetics Genomics 47, 37–47. https://doi.org/10.1016/j.jgg.2019.12.004

Zhou, L., Feng, Y., Wang, F., Dong, X., Jiang, L., Liu, C., Zhao, Q., Li, K., 2018. Generation of all-male-like sterile zebrafish by eliminating primordial germ cells at early development. Sci. Rep. 8, 1834. https://doi.org/10.1038/s41598-018-20039-3.

